# Activation and regulation of the dynein-dynactin-NuMA complex

**DOI:** 10.1101/2024.11.26.625568

**Authors:** Merve Aslan, Ennio A. d’Amico, Nathan H. Cho, Aryan Taheri, Juan M. Perez-Bertoldi, Yuanchang Zhao, Xinyue Zhong, Madeline Blaauw, Andrew P. Carter, Sophie Dumont, Ahmet Yildiz

**Author notes:** These authors contributed equally to this work (Merve Aslan, Ennio A. d’Amico, Nathan H. Cho).

## Abstract

During cell division, NuMA orchestrates the focusing of microtubule minus-ends in spindle poles and cortical force generation on astral microtubules by interacting with dynein motors, microtubules, and other cellular factors. Here, we used in vitro reconstitution, cryo-electron microscopy, and live cell imaging to understand the mechanism and regulation of NuMA. We determined the structure of the processive dynein/dynactin/NuMA complex (DDN) and showed that the NuMA N-terminus drives dynein motility in vitro and facilitates dynein-mediated transport in live cells. The C-terminus of NuMA directly binds to and suppresses the dynamics of the microtubule minus-end. Full-length NuMA is autoinhibited for its interactions with dynein and microtubules, but mitotically phosphorylated NuMA activates dynein in vitro and interphase cells. Together with dynein, activated full-length NuMA focuses microtubule minus-ends into aster-like structures. These results provide critical insights into the activation of NuMA and dynein for their mitotic functions.

## Introduction

Nuclear Mitotic Apparatus protein (NuMA) interacts with DNA, microtubules (MTs), motors, plasma membranes, importins, and other accessory proteins for fundamental cellular processes^1^. In interphase, NuMA localizes to the nucleus, where it organizes chromosomes and helps the formation of a single, round nucleus^2, 3^. After nuclear envelope breakdown, NuMA accumulates at MT minus-ends and recruits a minus-end-directed MT motor, dynein. Together with dynein, NuMA helps focus the minus-ends of the MTs into spindle poles and maintain a steady-state spindle structure^4–9^ (Fig. 1a). Minus-end-directed forces generated by dynein counteract forces generated by plus-end-directed kinesin-5s^10, 11^ to maintain spindle structure^12, 13^. NuMA also recruits dynein to the cell cortex^14–18^, where dynein anchors astral MTs and generates tension to determine the position and orientation of the spindle^18–20^. Consistent with its cellular roles, NuMA dysfunction causes defects in spindle mechanics and architecture, as well as micronuclei formation^1^.

**Fig. 1.**
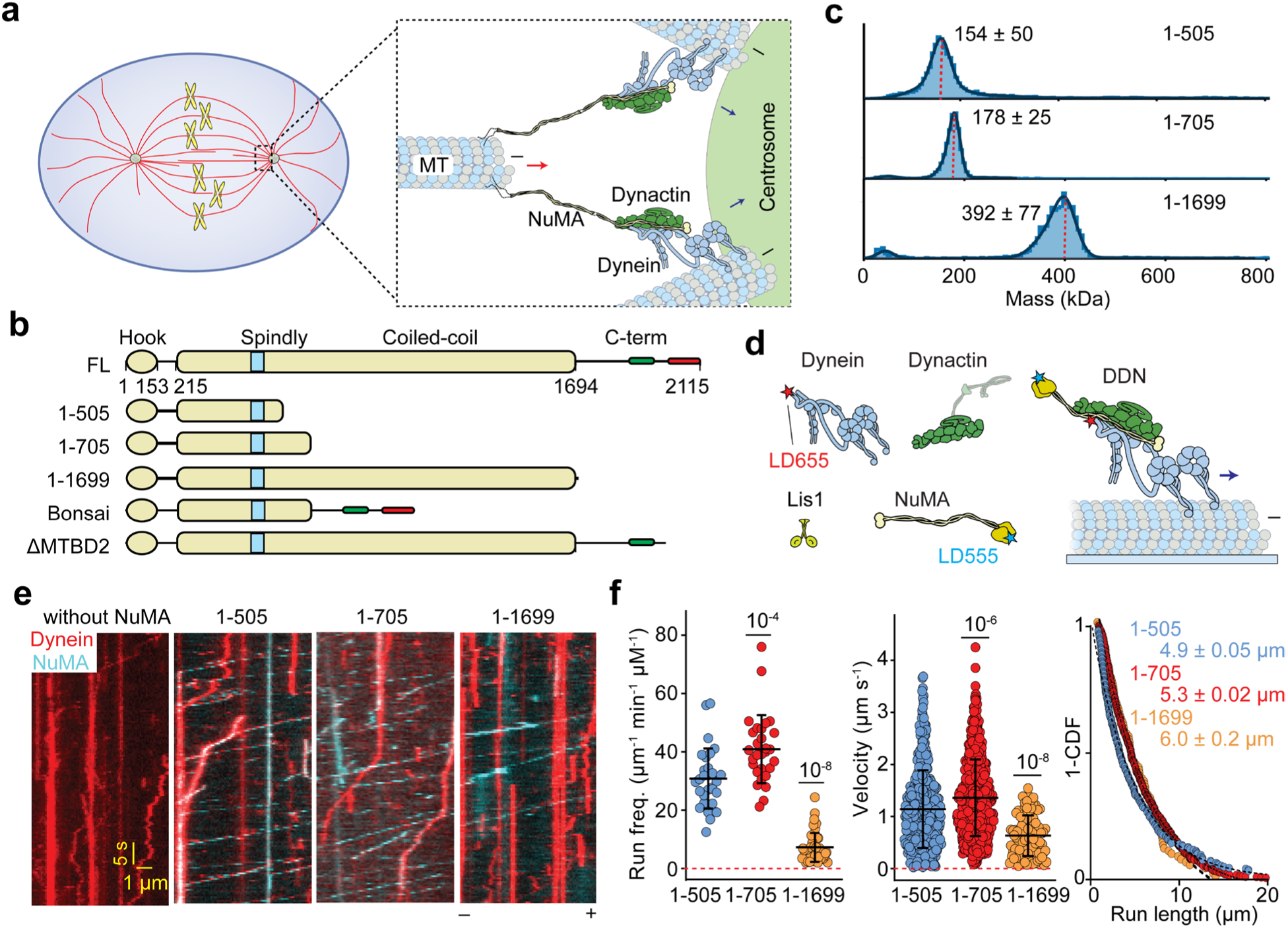
NuMA is a bona fide activator of dynein and dynactin. **a.** Schematic for the mitotic roles of NuMA at the spindle pole. NuMA recruits dynein/dynactin to the spindle poles, and together, they sort and focus the minus-ends of MTs. DDN complexes move toward the MT minus- end (blue arrows) while the NuMA interacts with the minus-ends of MTs as cargo. DDN motility moves the minus-ends of MTs towards the poles (red arrow). **b.** Domain organization and truncations of human NuMA. The Hook domain and the N-terminus of the long coiled-coil recruit dynein and dynactin. The C-terminus is disordered and interacts with MTs and other associated proteins. **c.** Mass photometry shows that NuMA 1-505, 1-705, and 1-1699 form homodimers (mean ± s.d.). Red dashed lines show the expected molecular weights of the dimeric forms of these constructs. **d.** Schematic depiction of the *in vitro* reconstitution and single-molecule imaging of DDN motility on surface-immobilized MTs. Lis1 was added to stimulate the formation of the DDN complex. Dynein and NuMA were labeled with LD655 and LD555 dyes, respectively. **e.** Kymographs show the processive motility of DDN complexes formed with different NuMA constructs *in vitro*. The assays were performed in 1 mM ATP and 100 mM KAc. **f.** The run frequency, velocity, and run length of single DDN complexes. The centerline and whiskers represent mean and s.d., respectively (from left to right, n = 29, 29, and 43 MTs for the run frequency, 418, 682, and 93 motors for the velocity and run length measurements). *P* values are calculated from a two-tailed t-test. The inverse cumulative distribution functions (1-CDF) of motor run length were fit to a single exponential decay to determine the mean run length (±s.e.).

NuMA is a large (237 kDa) and extended protein containing an N-terminal HOOK domain (amino acids (a.a.) 1-153) and a C-terminal region (a.a. 1700-2115) separated by a ∼210 nm long coiled-coil domain^21, 22^. The central coiled-coil facilitates dimerization^22^, provides the extension needed for properly crosslinking MTs and DNA^3, 23, 24^, as well as efficiently capturing astral MTs at the cell cortex^25^. The coiled-coil also prevents the binding of NuMA to chromosomes during mitosis^3^. The C-terminal region is disordered and contains two MT-binding sites (MTBD1 (a.a. 1914-1985) and MTBD2 (a.a. 2002-2115)), sites that bind to DNA and other associated factors, a clustering motif, and a nuclear localization signal^1^. MTBD2 can bind to the MT lattice^26^ and is required for spindle-pulling activity at the cortex in human cells^18, 19^. MTBD1 has a weaker affinity for MTs^27^, but it is required for spindle pole focusing^28, 29^ and accumulation of NuMA to the minus-end of MTs in mammalian cells^9^. There are contradictory reports for the distinct roles of MTBD1 and MTBD2 in spindle pole focusing and orientation in other cell types^30^, and it remains unclear how these MTBDs bind MTs and regulate MT dynamics.

Dynein forms a 1.4 MDa complex consisting of two copies of six subunits (heavy, intermediate, and light intermediate chains, and three light chains, Fig.1a)^31^. The dynein heavy chain is the largest subunit, which contains AAA+ ATPase and MT binding domains. The tail of the heavy chain facilitates homodimerization and recruitment of other subunits. Isolated dynein is autoinhibited and does not actively move along the MT^32, 33^. Activation of dynein motility requires recruitment to its cofactor dynactin (a 23-subunit, 1.2 MDa complex) via the coiled-coil region of adaptor proteins^34–36^. These activating adaptors contain several conserved motifs such as a specific binding site (i.e., CC1-box, HOOK, and EF-hand domains) for the C terminus of the dynein light intermediate chain (DLIC), a heavy chain binding site 1 (HBS1) motif for interaction with the dynein heavy chain (DHC) and a Spindly motif for interacting with the pointed end subcomplex of dynactin^37^. Studies in live cells showed that the N-terminal portion of NuMA (a.a. 1-505) is required for the recruitment of dynein to the cell cortex^18^. This region of NuMA contains a coiled coil and two regions reported to interact with the DLIC (an N-terminal HOOK domain and a CC1-box-like motif (a.a. 360–385))^38^, along with a predicted Spindly motif (a.a. 417–422)^18^ (Fig. 1b). Collectively, these observations suggest that the N-terminus of NuMA can form a complex with dynein and its cofactor dynactin. However, the presence of two distinct DLIC-binding sites, which has not been observed in any other activating adaptor, as well as the absence of a clear DHC binding site (HBS1), limits our understanding of how NuMA interacts with dynein and dynactin. It also remains unclear whether NuMA requires additional factors or post-translational modifications to activate dynein.

Our understanding of how NuMA orchestrates spindle assembly, maintenance, and orientation mostly comes from studies in cells. The molecular mechanisms by which NuMA is regulated and repurposed for its mitotic functions, recruits dynein/dynactin, localizes to the minus-ends of MTs, and focuses MT minus-ends remain unclear. The lack of in-depth *in vitro* biochemical and biophysical characterization of NuMA was partly due to its large size, low solubility, and tendency to form large clusters^18, 22, 39^. In this study, we used biochemical reconstitution, cryo-electron microscopy (cryo-EM), *in vitro* motility assays, and live cell imaging to investigate how human NuMA binds MTs and focuses MTs together with dynein. We demonstrated that the NuMA N-terminus forms a ternary complex with dynein and dynactin and activates dynein motility, whereas the C-terminal region preferentially binds and suppresses the dynamicity of the minus-end of MTs. Full-length (FL) NuMA only weakly binds MTs or activates dynein, whereas mitotically phosphorylated NuMA activates dynein motility *in vitro* and triggers dynein-driven transport in interphase cells. Together with dynein, active FL NuMA focuses minus-ends of MTs into aster-like structures *in vitro*. These results provide key insights into how NuMA organizes MTs and drives mitotic functions of dynein in the spindle body and at the cell cortex.

## Results

### The NuMA N-terminus assembles a DDN complex and activates dynein motility

To test the ability of NuMA to activate dynein motility *in vitro*, we recombinantly expressed and purified three human NuMA constructs containing the N-terminal coiled-coil (a.a. 1-505, 1-705, and 1-1699) from insect cells (Fig. 1b, Extended Data Fig. 1). Mass photometry showed that all three constructs form homodimers (Fig. 1c)^40, 41^. We next performed multi-color single-molecule motility assays on surface-immobilized taxol-stabilized MTs in the presence of dynein, dynactin, and Lis1, which favors the assembly of dynein with dynactin and adaptor protein (Fig. 1d)^42, 43^. In the absence of NuMA, dynein did not exhibit any processive runs on MTs. The addition of NuMA N-terminal constructs resulted in the activation of dynein motility and comigration of NuMA with dynein towards the minus-ends of MTs (Fig. 1e, Supplementary Movie 1). The velocity, run length, and run frequency of dynein/dynactin/NuMA (DDN) complexes with shorter N-terminal coiled coils were comparable to dynein-dynactin complexes with other activating adaptors, such as BICDR1^44^ (Fig. 1f). However, DDN with NuMA 1-1699 had ∼5-fold lower run frequency and ∼30% lower velocity than that of the shorter NuMA fragments, suggesting that the long coiled-coil has an inhibitory effect on DDN motility. These results showed that NuMA is a bona fide cargo adaptor that activates dynein-dynactin motility.

### Cryo-EM imaging of the DDN complex on MTs

To understand how NuMA forms a ternary complex with dynein and dynactin, we prepared DDN complexes with NuMA 1-705 in the presence of Lis1 and used a previously established cryo-EM workflow^45, 46^. We obtained a ∼6.5 Å resolution map of the part of the complex that contains dynactin, dynein tails, and the coiled-coil of NuMA (Fig. 2; Extended Data Figs. 2 and 3). The dynein motor domains, p150 arm, and Lis1 were not resolved. NuMA coiled-coil segments run along the length of dynactin, extending to the pointed end, and recruit two dynein motors (dynein-A and -B, Fig. 2b). The AlphaFold2 (AF2) model of NuMA predicts a break in the first coiled-coil between CC1a (a.a. 212-269) and CC1b (a.a. 276-396, Fig. 2a) that agrees with the density in our map (Fig. 2b-c). Our map also shows extra density visible outside the pointed end. Based on a high-confidence AF2 model of the interaction between NuMA and the dynactin pointed end (Extended Data Fig. 3a-b), we assigned this density to the Spindly motif site at the N-terminus of CC2 (408-705; Fig. 2b).

**Fig. 2.**
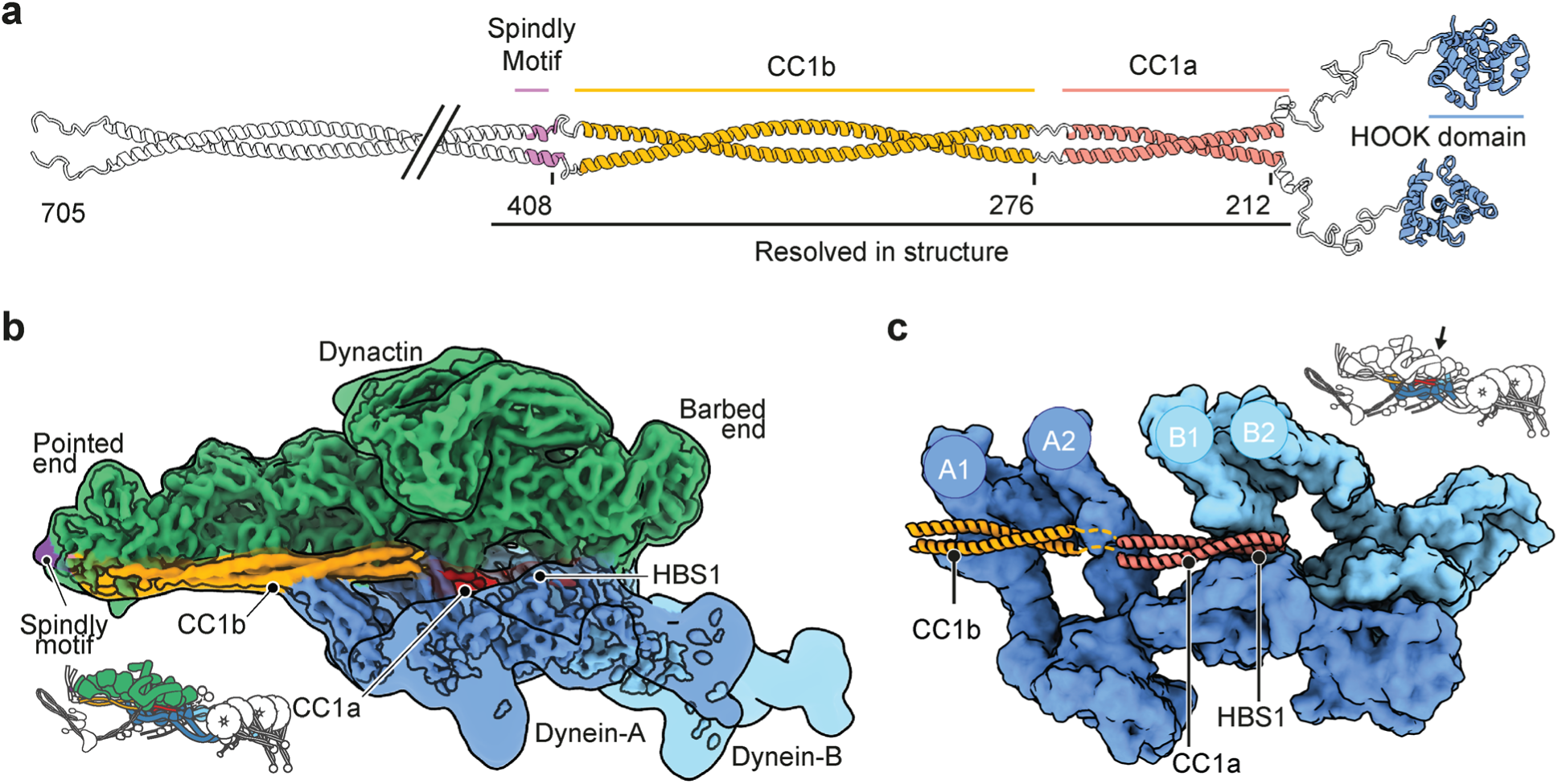
Cryo-EM structure of the DDN complex. **a.** Linearized AF2 prediction of NuMA 1-705 shown in the C-to-N direction. Only the CC1a, CC1b, and the Spindly motif are resolved. **b.** EM density map of dynein-dynactin bound to NuMA and Lis1 on MTs. The coiled-coil of NuMA runs along dynactin and recruits two dyneins to dynactin. The schematic on the bottom left shows the structure of dynein-dynactin-adaptor complexes on MTs. The dynein motor domains, p150 arm, and Lis1 are not resolved. **c.** The organization of the dynein-A and dynein-B interactions with the NuMA coiled-coil, with the dynactin subunits hidden. The arrow in the insert shows the viewing angle of the DDN complex. A1-A2 and B1-B2 are heavy chains of Dynein-A and Dynein-B shown in b.

In our model, CC1a contacts dynein-A at a site that includes DHC residues Y827 and R759. In other dynein/dynactin complexes^45, 46^, these residues bind the adaptor HBS1 motif, which consists of a conserved Gln residue and acidic patch (Fig. 3a-b). AF2 is unable to predict a NuMA HBS1- DHC interaction. However, from the position of the CC1a fragment, NuMA is likely to bind dynein-A using an acidic patch (residues 231-240) and a conserved region near Q215 (Fig. 3b). Although the HBS1 site is typically located in the middle of the coiled-coil in other activating adaptors^46^, this putative HBS1 site of NuMA is at the very N-terminus of the coiled-coil marked by a conserved proline residue (P212).

**Fig. 3.**
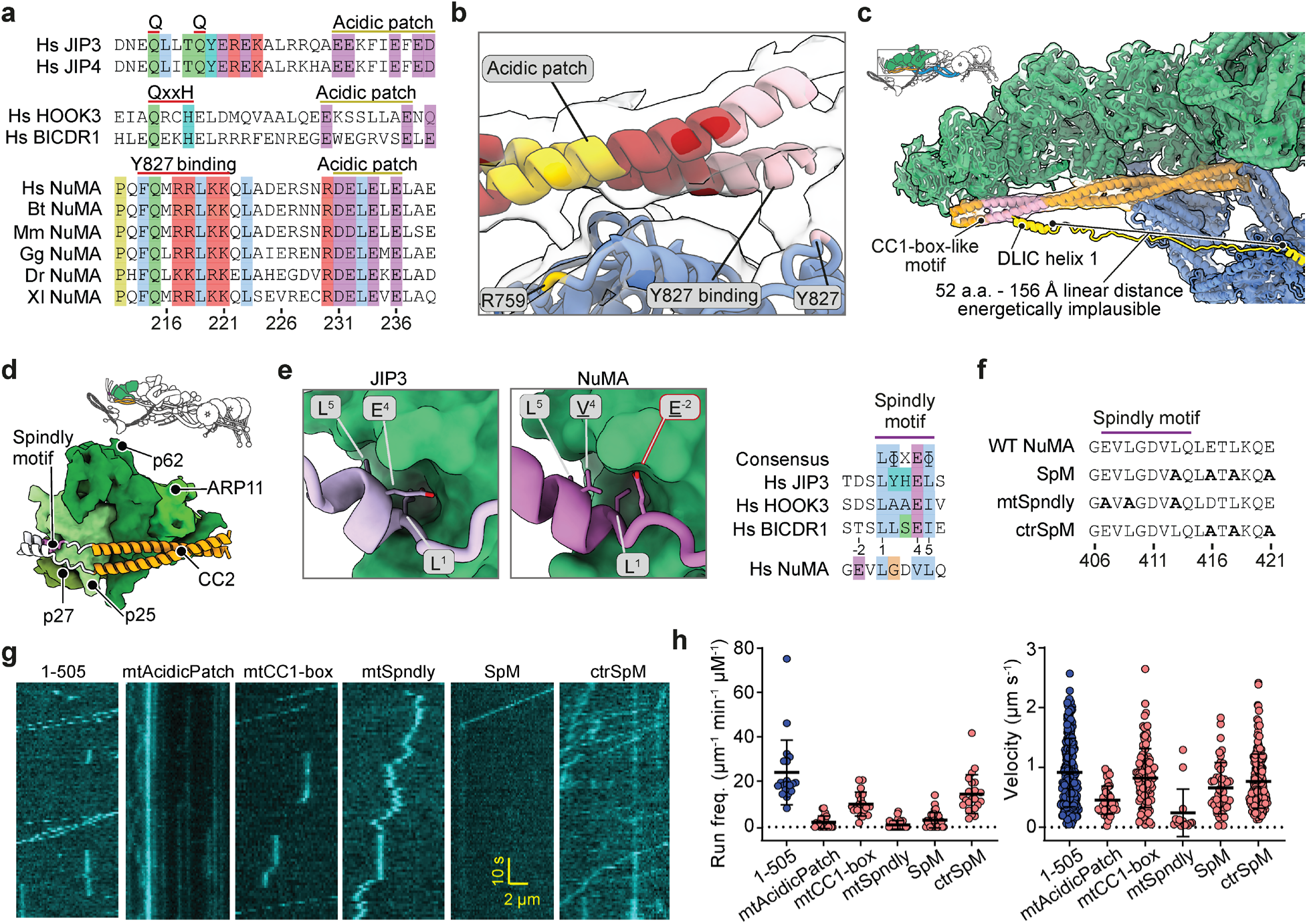
NuMA engages dynein–dynactin through a non-canonical Spindly motif, but not a CC1-box. **a.** Multiple sequence alignment of HBS1 from NuMA orthologs. Other activating adaptors with known HBS1 sequences are shown for comparison. **b.** Cryo-EM density and model of the NuMA HBS1-DHC interfaces. **c.** Close-up view of the CC1-box-like motif (pink) on NuMA in our EM density. A model of the DLIC tail with the unstructured region extended illustrates that the interaction is likely unfavored due to the distance. **d.** AF2 model of the NuMA interaction with the dynactin pointed end. Individual dynactin subunits are labeled. **e.** Spindly motif interactions with the dynactin pointed end in JIP3 (PDB: 8PR4) and NuMA (AF2, this study) illustrate the different positions of the interacting residues, as shown in a multiple sequence alignment of the Spindly motif in the adaptors JIP3, HOOK3, BICDR1, and NuMA. **f.** The design of the mutants introduced to the Spindly motif of NuMA. **g.** Example kymographs show the motility of DDN complexes assembled with WT and mutant NuMA 1-505 constructs. The assays were performed in 1 mM ATP and 100 mM KAc. **h.** The run frequency and velocity of single DDN complexes. The centerline and whiskers represent mean and s.d., respectively (from left to right, n = 21, 22, 19, 27, 31, and 22 MTs for the run frequency and 254, 42, 104, 14, 44, and 199 motors for velocity measurements).

NuMA is reported to bind the C-terminus of DLIC both through its Hook domain and a CC1-box- like motif^38^. In our structure, the proposed CC1-box-like motif sits on the far end of CC1b, in contact with the dynactin pointed end. Distance constraints and the lack of any additional density corresponding to the DLIC indicate that the CC1-box-like motif is unlikely to be engaged in interactions with dynein in our structure (Fig. 3c). While we cannot resolve the NuMA Hook domain due to its flexibility, an AF2 prediction suggests the DLIC1 helix, with its two conserved phenylalanines fits between the α8 helix and the α7-α8 loop of the NuMA Hook domain^38^ (Extended Data Fig. 3c-d). This position is consistent with previous mutational analysis^38^. Compared to the structure of the HOOK3 Hook domain – DLIC1 helix complex^47^, the NuMA Hook domain lacks an α7 helix, which translates to closer contact with the α8 helix (Extended Data Fig. 3c).

NuMA makes multiple contacts with the pointed end subunits of dynactin through CC1b and the Spindly motif on CC2 (Fig. 3d), but its Spindly motif differs from the consensus LxxEΦ motif (where Φ is hydrophobic) found in other adaptors^46^. In BICDR1 and JIP3, the hydrophobic residues in this motif bind a pocket on the pointed-end p25 subunit, while the glutamate shields the pocket and interacts with N20 on p25 (Fig. 3e). In NuMA, the model predicts the Spindly motif sequence is LGDVL. The two leucines (L409 and L413) bind the hydrophobic pocket on p25, whereas a glutamate outside the core motif (E407) contacts N20 (Fig. 3e; Extended Data Fig. 3e). Collectively, our structural analysis showed that NuMA binds dynein and dynactin through an HBS1 site and a previously unknown Spindly motif, and that a proposed CC1-box-like motif is not likely involved in complex formation.

To validate our structural model, we introduced point mutations to the acidic patch, proposed CC1-box-like motif, and the Spindly motif of NuMA 1-505, and tested how these modifications affect DDN motility *in vitro* (Fig. 3f-h). Consistent with our model, reversing the charges of the critical residues in the acidic patch (<u>DE</u>L<u>E</u>L<u>E</u> → <u>KR</u>L<u>R</u>L<u>R</u>, Fig. 3a) nearly fully eliminated dynein motility, while previously proposed mutations to the CC1-box-like motif (A363V/A369V)^38^ had only a minor effect. Alanine mutations to the critical residues within the Spindly motif that our model predicts (mtSpndly) nearly eliminated dynein motility. A previously suggested position for the Spindly motif^18^ is downstream from the predicted sequence but overlaps with it at residue L413. Alanine substitutions to the previously reported Spindly mutant (SpM) also impaired dynein activity both in cellular assays^18^ and in our *in vitro* assays (Fig. 3f-h, Extended Data Fig. 3f). Reverting the residue 413 back to leucine (L) in SpM (ctrSpM) nearly fully rescued DDN motility, showing that SpM disrupts dynein motility due to the mutation to the Spindly motif at L413. These results strongly support our structural model and are consistent with the requirement of the Spindly motif in the activation of dynein motility by other adaptor proteins^48, 49^.

### MTBD1 Preferentially Binds the Minus-end of HMTs

To understand the interaction of the NuMA C-terminus with MTs, we purified NuMA C (a.a. 1700-2115), MTBD1 (a.a. 1811 – 1985; previously referred to as NuMA TIP^30^), and MTBD2 (a.a. 2002 – 2115; Fig. 4a, Extended Data Fig. 4a-b). Although earlier studies referred to the NuMA C-terminus as a globular domain^50, 51^, AF2 predicts that it is almost fully disordered (Extended Data Fig. 4c), consistent with the interaction of this region with a diverse array of cellular factors and its role in liquid-liquid phase separation of NuMA both *in vitro* and *in vivo*^39^. The C-terminal constructs lack the coiled-coil region and primarily appeared as monomers in mass photometry (Fig. 4b). We found that NuMA C, which contains both MTBD1 and MTBD2 sites, appeared both on the lattice and the tip of MTs (Fig. 4c, Extended Data Fig. 5a). Consistent with a previous report^27^, MTBD1 weakly interacted with the MT lattice. However, it mostly appeared as bright puncta at the tip of a subset of MTs. In comparison, MTBD2 densely decorated the MT lattice, albeit at a lower degree than NuMA C (Fig. 4c). NuMA constructs that lack the C-term failed to bind MTs (Extended Data Fig. 5b).

**Fig. 4.**
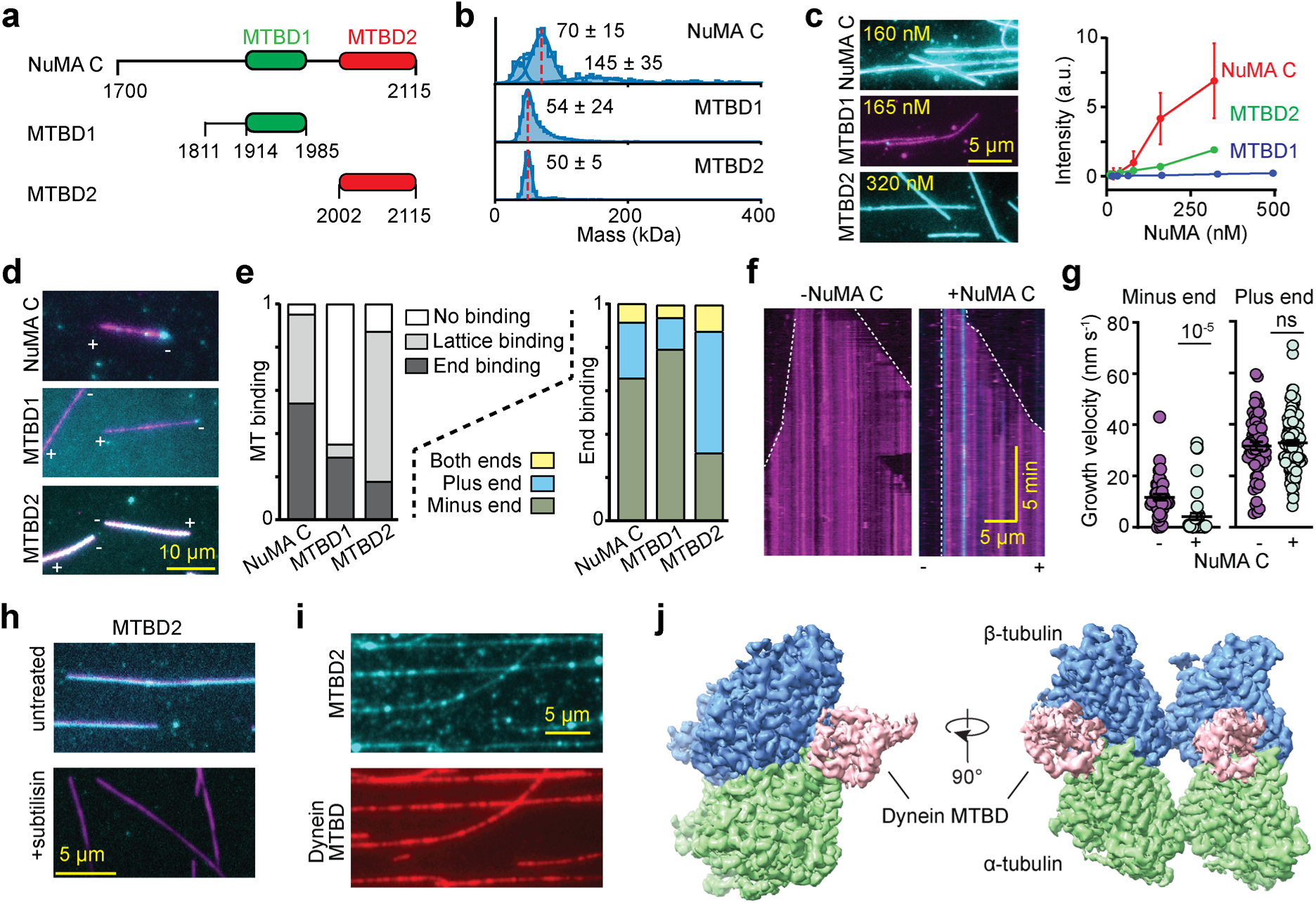
The C-terminus of NuMA preferentially binds to the minus-end of MTs. **a.** The NuMA C-terminal region (NuMA C) contains two MTBDs (MTBD1 and 2). **b.** Mass photometry shows that the C-terminal constructs primarily form monomers (mean ±s.d.). Red dashed lines show the expected molecular weights of the monomers. **c.** (Left) Example pictures show NuMA C and MTBD2, but not MTBD1, densely decorated surface-immobilized MTs. (Right) The fluorescence intensity of NuMA constructs per MT length (from left to right, n = 282, 324, 328, 320, 316, 348 for NuMA C, 296, 195, 310, 313, 300, 318 for MTBD1, and 254, 200, 201, 206, 200, and 161 for MTBD2). The centerline and whiskers represent mean and s.d., respectively. **d.** Example pictures show MT binding of 25 nM NuMA C, 165 nM MTBD1, and 4 nM MTBD2. MT polarity is determined from the directionality of plus-end-directed Cy5-labeled K490 motors (not shown). **e.** Normalized MT binding and end binding preference of C-terminal constructs (from left to right, n = 107, 117, and 142 MTs). **f.** Kymographs of dynamic MTs with GMPCPP MT seeds in the presence and absence of 100 nM NuMA C. Minus-end accumulation of NuMA C stops its growth and shrinkage. **g.** Growth velocities of the MT plus and minus ends in the absence and presence of NuMA C (from left to right, n = 41, 43, 59, and 126 MT growth events). The centerline and whiskers represent mean and s.e.m., respectively. *P* values are calculated from a two-tailed t-test. **h.** MTBD2 exhibits little to no binding to subtilisin-treated MTs. **i.** MTs are co-decorated with 50 nM MTBD2 (cyan) and 500 nM dynein-MTBD (red). **j.** Cryo-EM reconstruction of the co-decorated MTs only shows density corresponding to dynein-MTBD. No additional density that could correspond to NuMA was observed. In c, d, f, and h, NuMA and MTs are colored in cyan and magenta, respectively.

Previous studies in live cells hinted that MTBD1 is sufficient for binding of NuMA to minus-ends of MTs, whereas there were conflicting reports on whether MTBD1 or MTBD2 prefers curled ends of MTs and is required for localization of NuMA at the plus-end tip of astral MTs^9, 26, 30, 52^. We quantified the lattice and tip binding preference of NuMA C, MTBD1, and MTBD2 at sub-saturation concentrations (Fig. 4d-e, Extended Data Fig. 5c). MTBD1 rarely binds to the lattice and strongly (∼80%) prefers binding to the minus-end when it lands on the MT. MTBD2 decorates the lattice even at low concentrations and does not exhibit strong tip binding (∼20%), but it prefers to land on the plus-end when it only appears at the MT end. NuMA C exhibits the MT binding features of both MTBD1 and MTBD2, appearing both at the ends and the lattice of MTs, but prefers the minus-end when it only appears at the tip of the MT (Fig. 4d-e). These results show that NuMA can recognize and bind to the minus-ends of MTs through MTBD1^9^, but it also interacts with the lattice and the plus-end through MTBD2^26, 52^.

To determine whether MT binding of NuMA regulates MT dynamics, we immobilized MT seeds stabilized with a nonhydrolyzable GTP analog, GMP-CPP, to the glass surface and monitored active polymerization and depolymerization of MTs in the presence of free tubulin, GTP, and NuMA C (Fig. 4f). We observed NuMA C binding to both ends and the lattice of dynamic MTs. The accumulation of NuMA C at the MT minus-end almost completely stopped its growth, whereas there was little to no effect on the polymerization dynamics at the plus-end (Fig. 4g). These results show that the C-terminal region of NuMA caps and stabilizes the minus-ends of MTs.

We next sought to understand how MTBD2 binds to the MT lattice. The MT association of NuMA C weakened upon increasing the salt concentration to 200 mM KAc (Extended Data Fig. 5d), suggesting electrostatic interactions between NuMA and MTs. The interaction of NuMA C and MTBD2 with MTs was also substantially reduced when the tubulin tails were cleaved by a serine protease, subtilisin, indicating that tubulin tails play an important role in MT binding of NuMA (Fig. 4h, Extended Data Fig. 5e-f). To further test whether MTBD2 binds to the globular fold or the flexible tails of polymerized tubulin, we imaged MTs decorated with MTBD2 at near saturation. To reliably distinguish α and β tubulin and determine the MT seam, we co-decorated MTs with a construct that contains the dynein MTBD in a high-affinity state (dynein-MTBD)^53^ (Fig. 4i) and imaged these MTs using cryo-EM (Extended Data Figs. 6 and 7). The pseudo-helical reconstruction and symmetry expansion of MTs at 3.3 Å resolution revealed a clear density that corresponds to the dynein-MTBD^53^. However, we did not observe any additional density that could correspond to NuMA MTBD2 (Fig. 4j; Extended Data Fig. 6), even though NuMA MTBD2 densely localizes to MTs under these conditions (Fig. 4j, Extended Data Figs. 6 and 7c). These results support that MTBD2 of NuMA interacts with the flexible tails of tubulin, which cannot be directly observed in our cryo-EM reconstruction.

### NuMA is Rescued from Autoinhibition by Phosphorylation of its C-terminus

We turned our attention to full-length (FL) NuMA, which contains both dynein and MT interaction sites (Extended Data Fig. 8a). Previous studies, which used denaturation/renaturation of NuMA FL and rotary shadow imaging, reported that NuMA forms oligomers with 6-12 long flexible arms^19,38^. We imaged NuMA FL without protein denaturation using negative stain EM. Consistent with these reports, NuMA FL formed clusters that project ∼200 nm long and flexible extension arms (Extended Data Fig. 8b-c). In contrast, NuMA 1-1699 only forms ∼200 nm long extension arms without clusters (Extended Data Fig. 8b), demonstrating that the C-terminus is responsible for the clustering of NuMA FL, and long extension arms correspond to the coiled-coil domain. Compared to these reports^19, 38^, we observed NuMA forming larger clusters that project a highly variable number of flexible arms. These discrepancies may be due to the differences in sample preparation. Clustering of NuMA FL was also observed in fluorescence imaging assays (Fig. 5a). These observations are in line with the punctate signals of NuMA at the cell cortex^18^ and the proposed function of NuMA in crosslinking and focusing MT minus-ends into larger asters in acentrosomal spindles^52^.

**Fig. 5.**
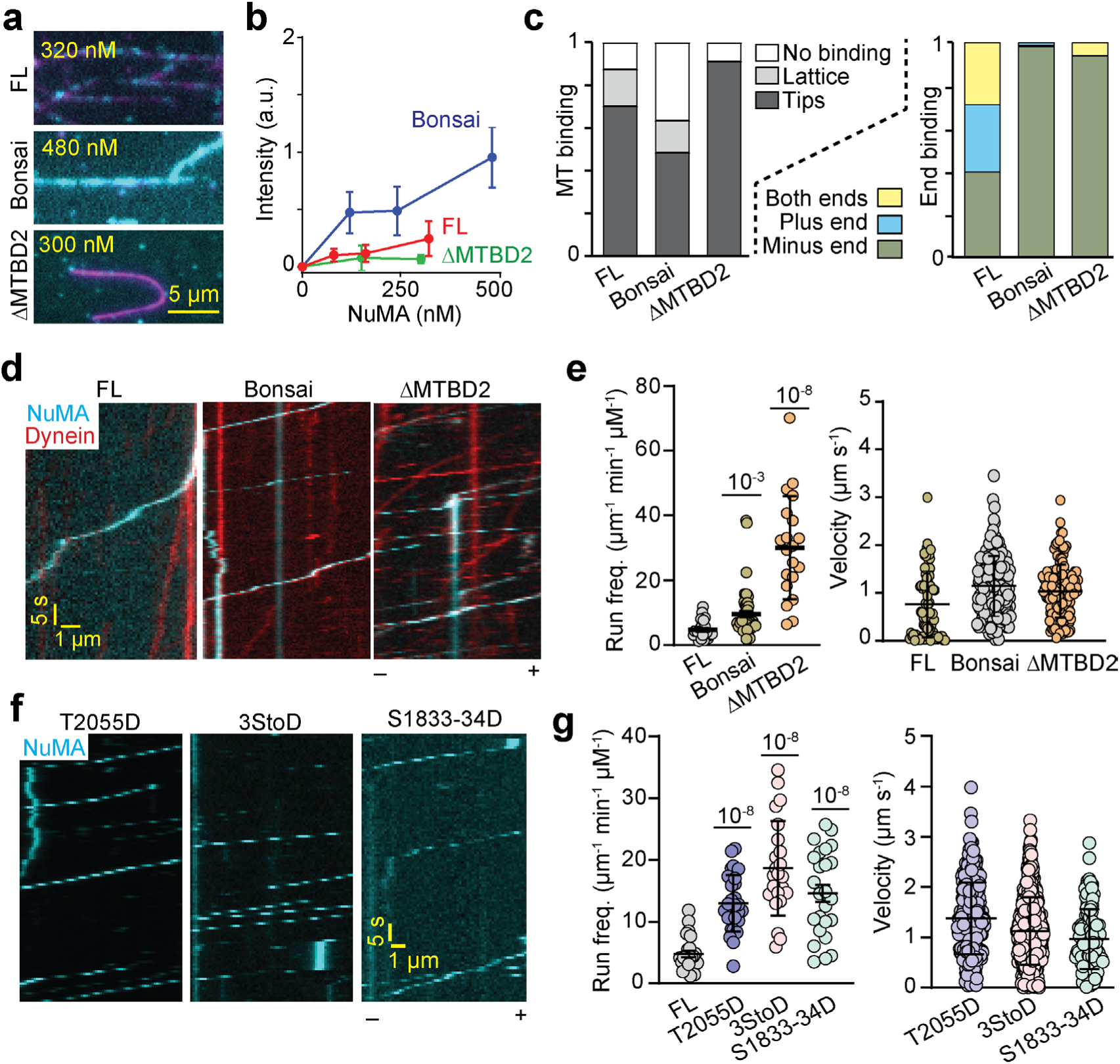
NuMA-dynein interaction is activated by the phosphorylation of its C-terminus. **a.** Example pictures show MT (magenta) binding of NuMA FL, Bonsai, and ΔMTBD2 (cyan) under different concentrations. **b**. The intensity of NuMA constructs fused to mNeonGreen (mNG) per MT length (from left to right, n = 30, 203, 203, 348 MTs for NuMA FL; 30, 251, 237, 276, 237 for Bonsai; and 30, 48, 68 MTs for ΔMTBD2). Error bars represent s.d. **c.** Normalized MT binding and end binding preference of 150 nM NuMA FL, 120 nM Bonsai, and 300 nM ΔMTBD2 (n = 104,126, and 91 MTs, from left to right). **d**. Kymographs show processive motility of DDN complexes assembled with NuMA FL, Bonsai, and ΔMTBD2. **e**. The run frequency and velocity of DDN complexes assembled with NuMA FL, Bonsai, and ΔMTBD2 (from left to right, n = 24, 52, and 20 MTs for the run frequency and 85, 154, and 152 motors for the velocity measurements). **f.** Kymographs show the processive motility of DDN complexes assembled with phosphomimetic mutants of NuMA FL. The NuMA signal appears discontinuous due to time sharing with other fluorescent channels (not shown). **g.** The run frequency and velocity of DDN complexes assembled with phosphomimetic mutants of NuMA FL (from left to right, n = 24, 28, 23, and 25 MTs for run frequency and 286, 435, and 136 motors for velocity measurements). In e and g, the centerline and whiskers represent mean and s.d., respectively. *P* values are calculated from a two-tailed t-test.

We next tested how NuMA FL interacts with dynein and MTs. Unlike NuMA C, NuMA FL had a low affinity for MTs and exhibited no clear preference for tip binding (Fig. 5a-c). NuMA FL also resulted in ∼10-fold less frequent processive runs and ∼2-fold slower movement than complexes assembled with NuMA 1-705 under the same conditions (Figs. 1e-f and 5d-e, Extended Data Fig. 8d). These results indicate that NuMA FL is autoinhibited for its interactions with dynein and MTs. NuMA autoinhibition may involve interactions between the N- and C-terminal regions mediated by the central coiled-coil, analogous to other cargo adaptors of dynein^54–56^. We tested this possibility by truncating the central coiled-coil and the extreme C-terminus of NuMA. NuMA Bonsai, which contains both the N-terminal (1-705) and C-terminal (1700-2115) region, but lacks the intervening coiled-coil domain^3, 18^, binds to MTs more efficiently than NuMA FL, but less efficiently than NuMA C (Fig. 5a-b, Extended Data Fig. 8e). It prefers binding to the minus-end of MTs (Fig. 5c). NuMA Bonsai also resulted in more frequent dynein runs than NuMA FL, albeit at a lower frequency than NuMA 1-705 (Figs. 1e-f and 5d-e). A NuMA construct that lacks MTBD2 (ΔMTBD2) very weakly interacted with MTs and preferred to bind the MT minus-end (Fig. 5c), consistent with MTBD2 being the main driver of MT binding, and MTBD1 being the site of minus-end recognition of MTs. Unlike NuMA Bonsai, ΔMTBD2 stimulated dynein motility nearly as efficiently as 1-705 (Fig. 5e). Collectively, these results indicate that the central coiled-coil is involved in the autoinhibition of dynein binding at the N-terminus and MT binding at the C-terminus. The removal of this region results in partial activation of NuMA, suggesting that the N- and C-terminal regions are also part of the autoinhibitory mechanism, such that the presence of the C-terminus prevents full activation of dynein at the N-terminus and the N-terminus reduces MT binding of the C-terminus. We also note that DDN Bonsai and ΔMTBD2 moved at a similar velocity to that of complexes assembled with NuMA 1-705 (Figs. 1e-f and 5e), suggesting that the presence of NuMA MTBDs does not cause intermittent pausing or dragging against dynein motility.

We reasoned that NuMA may be rescued from autoinhibition and activated for its mitotic functions through phosphorylation^57, 58^. Studies in live cells revealed that, after nuclear envelope breakdown, NuMA is phosphorylated by CDK1 at T2055 to enhance its MT binding and inhibit its localization to the cell cortex^15, 59^. S1969, S1991, and S2047 phosphorylation by Aurora-A kinase facilitates the translocation of NuMA from spindle poles to the cell cortex^60^. NuMA is also phosphorylated by Plk1 at S1833/34 ^61^, which increases the rate of NuMA turnover at spindle poles and the cell cortex^62^. We generated phosphomimetic mutants of the Aurora A (S1969D, S1991D, and S2047D; 3StoD hereafter)^57, 60^, CDK1 (T2055D)^59^, and Plk1 (S18833/34D)^62^ phosphorylation sites and tested how these modifications affect dynein activation of NuMA FL (Extended Data Fig. 8a). Similar to wild-type NuMA FL, these mutants exhibited low affinity to bind MTs (Extended Data Fig. 9a-b). However, they exhibited differences in their end-binding preference. While T2055D was more likely to bind to the plus-end, S1833/34D was preferably localized to the MT minus end (Extended Data Fig. 9c). We concluded that phosphorylation of NuMA at these sites does not substantially affect its MT binding.

Unlike MT binding, all three phosphomimic mutants resulted in up to a 4-fold increase in the run frequency of DDN complexes (Fig. 5f-g, Supplementary Movie 2). The velocity of these complexes was comparable, but their run frequency was still lower than that of DDN complexes assembled with NuMA 1-705 (Fig. 5f-g, Extended Data Fig. 9d-e). These results indicate that interphase NuMA is autoinhibited for its interactions with dynein and partially activated by mitotic phosphorylation of its C-terminus by Aurora A, CDK1, and Plk1.

### Mitotically Phosphorylated NuMA Activates Dynein in Interphase Cells

To investigate whether NuMA can activate dynein in cells, we leveraged synthetic activation of dynein in peroxisome transport through chemically inducible dimerization of FRB and FKBP tags upon rapamycin addition (Fig. 6a)^63^. Peroxisomes typically diffuse freely through the interphase cytoplasm. We exogenously expressed a peroxisome marker (PEX3) fused with mEmerald and FKBP, and fused FRB to different NuMA constructs that lack the NLS motif. If the NuMA construct activates dynein-mediated transport, we expect PEX3-mEmerald to move to the centrosome, where minus-ends terminate, typically in the perinuclear region.

**Fig. 6.**
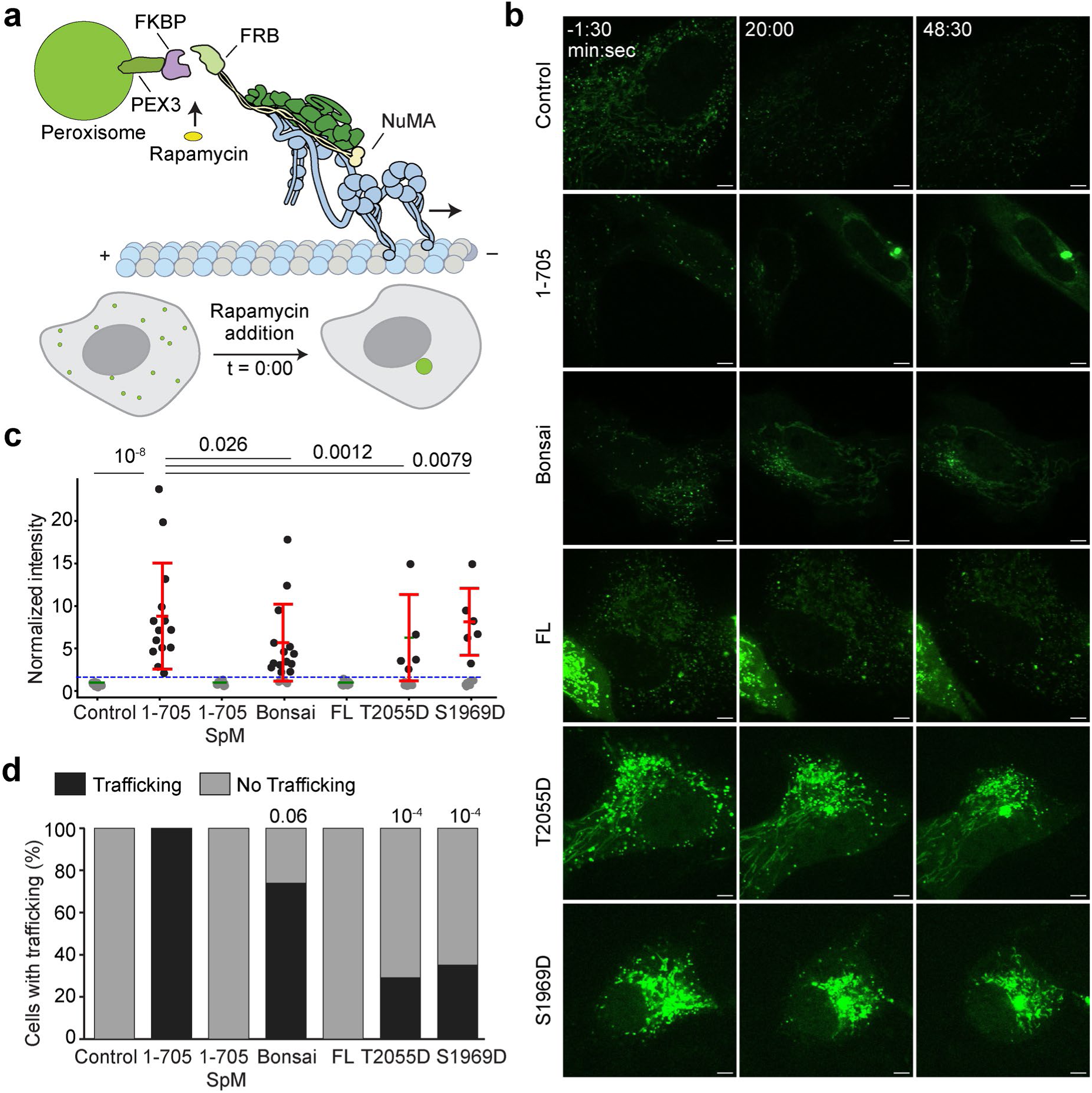
Mitotically-phosphorylated NuMA activates dynein in interphase cells. **a.** Cartoon depiction of the peroxisome trafficking assay. Rapamycin addition (time 0:00) induces recruitment of a NuMA construct to peroxisomes. If the NuMA construct activates dynein, these peroxisomes are trafficked towards the centrosome. **b.** Representative confocal images of peroxisomes (PEX3, green) before and after rapamycin addition in WT RPE1 cells (control) and RPE1 cells expressing different NuMA constructs. PEX3-mEmerald-FKBP levels vary between cells due to variations in transfections. Scale bar = 5 µm. **c.** The ratio of final (45 min after rapamycin addition) mEmerald intensity to initial mEmerald intensity at the centrosome. Cells were scored as trafficking (black) if the ratio was above the threshold (2, blue dashed line) or no trafficking (grey) if below the threshold (from left to right, n = 16, 14, 15, 19, 22, 17, and 17; two independent experiments). Error bars represent mean ±s.d. of trafficking cells. *P* values are calculated from a two-tailed t-test and Fisher’s exact test, respectively, in comparison to the 1-705 condition. **d.** Percent of cells with peroxisome trafficking, from the same samples as c.

In agreement with our *in vitro* results (Fig. 1), we found that NuMA 1-705 efficiently traffics peroxisomes to the centrosome in interphase cells (Fig. 6b-d, Extended Data Fig. 10, Supplementary Movie 3). A construct containing the previously predicted Spindly mutation (NuMA 1-705 SpM)^18^ fully eliminated trafficking, verifying that trafficking results specifically from NuMA-dynein interactions, as opposed to an indirect mechanism. NuMA-Bonsai, which additionally has NuMA’s C-terminus that can bind MTs, also trafficked peroxisomes. Notably, NuMA FL did not cause peroxisomes to cluster. In comparison, phosphomimetic NuMA FL constructs (S1969D and T2055D) trafficked peroxisomes, albeit in a lower percentage of cells compared to NuMA 1-705. Importantly, differences in NuMA expression level or pre-accumulation of NuMA to the centrosome do not explain the differences in trafficking between NuMA FL and phosphomimetic NuMA FL constructs (Extended Data Fig. 10c-d and 10g). These results are consistent with *in vitro* motility assays (Fig. 5f-g) and show that NuMA FL can activate dynein in interphase cells when mitotically phosphorylated. Therefore, these post-translational modifications on NuMA might not only control its cellular localization, but also enable it to form active DDN complexes with dynein.

### NuMA Focuses MTs into Asters Together with Dynein

We next investigated how NuMA affects the organization of MTs with or without dynein. To test whether NuMA can crosslink and bundle MTs, we enabled the free movement of MTs while keeping them near the glass surface using the depletion forces generated by a crowding agent (Fig. 7a). In the absence of NuMA, we observed little to no bundling of MTs. NuMA FL was bound to freely floating MTs but did not cause substantial MT bundling, probably due to its low MT affinity. Although NuMA C or MTBD2 lack the coiled-coil region and do not form a dimer, they bundled the MTs to a significant extent^24^ (Fig. 7b).

**Fig. 7.**
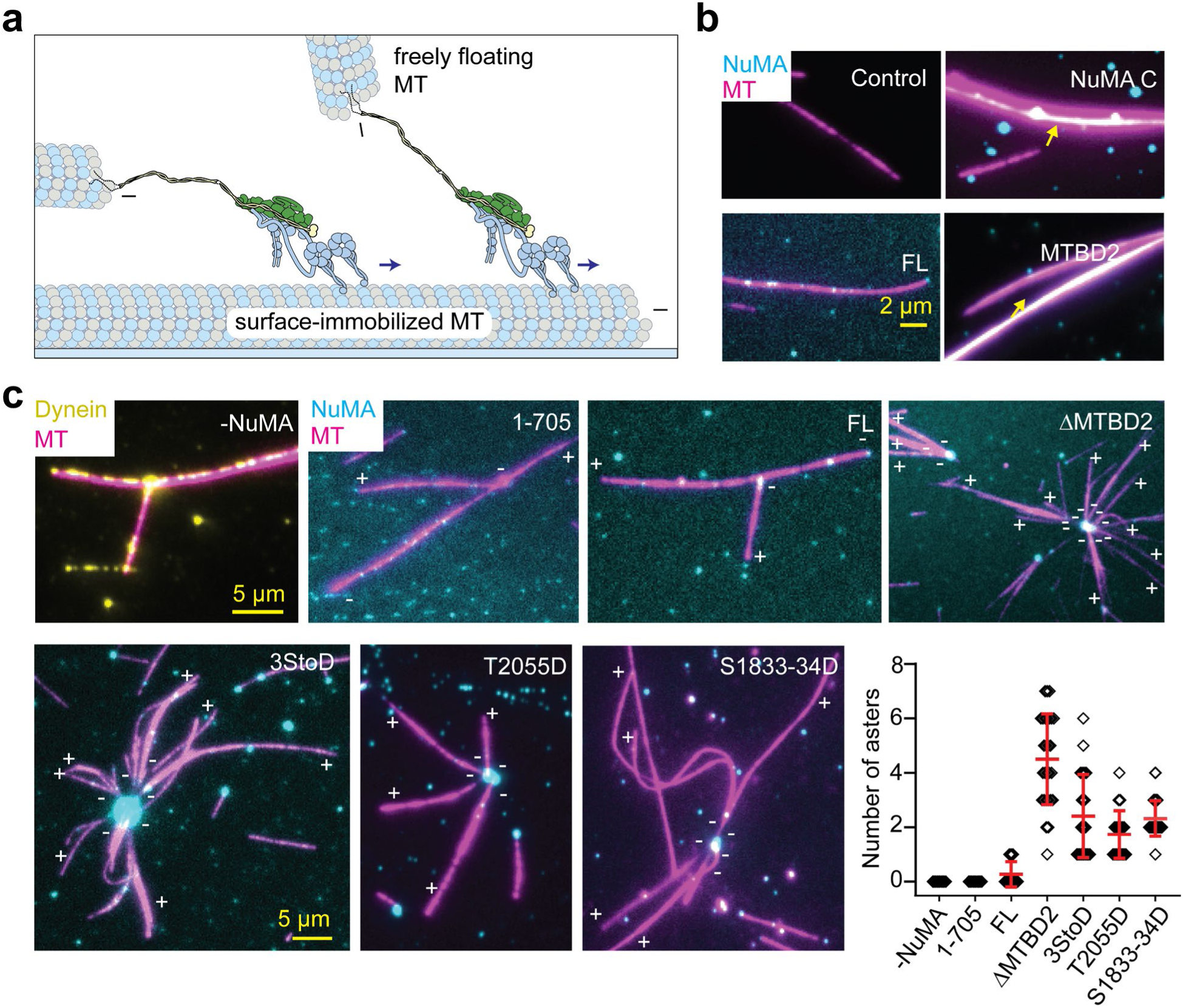
NuMA focuses MT minus-ends into asters with dynein/dynactin. **a.** Schematic shows alignment and focusing of the minus-end of the freely floating MTs to the minus end of a surface-immobilized MT by the DDN complex. **b.** Example pictures show that NuMA FL weakly interacts with freely diffusing MTs, whereas NuMA C and MTBD2 form MT bundles (yellow arrows). **c.** Example pictures show the organization of MT minus-ends into aster-like structures in the presence of NuMA, dynein, and dynactin. MT polarity is determined from the minus-end-directed motility of DDNs (not shown). The disappearance of the MT signal near the center of the asters in DDN 3StoD and T2055D conditions is due to the bending of MTs in the z direction near the NuMA cluster, which positions them away from the evanescent field of TIRF excitation. (Bottom right) The number of asters detected in an 80 µm x 80 µm imaging area in the presence of different NuMA constructs. Asters were defined as the coalescence of the minus ends of at least three MTs (from left to right, n = 10, 10, 11, 28, 24, 19, and 31 images). In b and c, NuMA, MTs, and dynein are colored in cyan, magenta, and yellow, respectively.

To test whether minus-end recognition and dynein activation of NuMA are sufficient to focus MTs into spindles, we mixed freely diffusing MTs and NuMA with or without dynein/dynactin and imaged their accumulation on surface-immobilized MTs (Fig. 7c, Extended Data Fig. 11). In the absence of dynein and dynactin, FL NuMA constructs decorated the MT surface and occasionally caused bundling and coalescence of MTs, but we did not observe the formation of aster-like MT organization (Extended Data Fig. 11a). Similar results were observed for dynein and dynactin in the absence of NuMA (Fig. 7c). Aster formation was also not observed when dynein and dynactin were mixed with either NuMA 1-705 that does not bind MTs or NuMA FL that is mostly autoinhibited and has very low dynein activation.

Remarkably, the addition of dynein, dynactin, and mitotically phosphorylated FL NuMA constructs resulted in a robust assembly of MTs into asters (Fig. 7c, Supplementary Movie 4). During aster formation, typically, NuMA first appears as bright puncta at the minus-end of MTs. The coalescence of these MTs with another MT results in its minus-end-directed transport by DDN. As a result, we observe a large accumulation of NuMA and focusing of MTs from their minus-ends at the center of the asters. Out of the constructs we tested, ΔMTBD2 was most efficient in forming aster-like structures together with dynein and dynactin (Fig. 7c), likely because removal of MTBD2 resulted in substantial activation of DDN motility (Fig. 5d) without affecting the recognition of the minus-ends of MTs through MTBD1 (Extended Data Fig. 8e). These results indicate that both the MT minus-end recognition and dynein activation of NuMA are essential for focusing of the minus-ends into aster-like structures.

## Discussion

We used biochemical reconstitution with purified components and single particle cryo-EM and light microscopy to directly observe how NuMA activates dynein motility, binds MTs, affects MT dynamics, and organizes the MT filaments together with dynein *in vitro*. We demonstrated that the N-terminus of NuMA forms a processive DDN complex with dynein and dynactin, triggering dynein-driven transport when targeted to intracellular cargos in live cells. A puzzling feature of NuMA activation of dynein was the presence of both a Hook domain and a CC1-box-like motif. Our findings suggest that the CC1-box-like motif does not engage with DLIC, implicating the Hook domain as the primary mediator of this interaction. While we cannot exclude that the CC1-box-like motif interacts with dynein in other cellular contexts, our observation that the Hook domain is the only one relevant for the assembly of the full DDN complex is consistent with cellular observations^38^. Our structure also indicates that NuMA binds dynein and dynactin through a unique Spindly motif and HBS1 sequences. These results shed light on the diverse mechanisms by which dynein and adaptors interact, suggesting a flexible, functionally specific framework for these interactions.

We also characterized the MT binding of MTBD1 and MTBD2 sites at the C-terminal region of NuMA. MTBD2 is the main driver of MT binding by interacting with the C-terminal tails of polymerized tubulin along the length of the MT. In comparison, MTBD1 weakly binds to the MT lattice and recognizes the minus-end of MTs. Minus-end localization of NuMA suppresses the dynamics of growth and shrinkage at this end. It remains to be determined which residues in MTBD1 and MTBD2 are responsible for MT binding. Future studies are also required to reveal whether MTBD1 recognizes the minus-end due to the exposed α-tubulin subunit or unique protofilament angles and curvature at the minus end^64, 65^.

We found that NuMA FL is autoinhibited for its interaction with dynein. The N-terminal Hook domain and the part of the coiled-coil (1-505 and 1-705) were most effective in the activation of dynein motility, whereas the inclusion of the rest of the coiled-coil (1-1699) disfavored the assembly of DDN complexes. The central coiled-coil may fold onto either of these domains for their inhibition, in line with our previous observation that it prevents NuMA from binding chromosomes at mitosis^3^. The ability of NuMA FL to interact with dynein was most significantly enhanced by the truncation of MTBD2, suggesting that the N-terminus is primarily inhibited by the NuMA C-terminus. Similarly, NuMA FL also weakly interacts with the MTs, suggesting that interactions between N- and C-termini may also block the MT interaction sites at the C-terminus. Future studies will be required to reveal the structure and mechanism of NuMA autoinhibition.

In interphase cells, NuMA is localized to the nucleus and is unable to decorate MTs or interact with dynein. Our peroxisome assays in live cells indicate that NuMA is inhibited from interacting with dynein even if it does not localize to the nucleus in interphase cells, providing molecular insights into the autoinhibition and activation of NuMA during mitosis. The autoinhibitory interactions may play a major role in repurposing NuMA for its diverse array of cellular functions. Our NuMA FL may represent NuMA’s interphase state, which does not strongly interact with MTs and dynein. In comparison, the assembly of NuMA with dynein and dynactin was partially activated by phosphorylation of its C-terminus by mitotic kinases Aurora A, CDK1, and Plk1. Full activation of dynein interactions may require other posttranslational modifications or interaction partners, such as RanGTP and Gαi^1^. Consistent with this view, a recent study reported that activation of dynein by NuMA requires CDK1 phosphorylation of the N-terminus of the coiled-coil^66^.

Our results indicate that mitotic phosphorylation of NuMA does not simply regulate NuMA localization, but it is required to activate motility and force generation of dynein for spindle pole focusing and cortical force generation. While the underlying mechanism remains unclear, these phosphorylation events may disrupt some of the autoinhibitory interactions between the N- and C-termini of NuMA and allow NuMA to attain an open conformation to interact with its partners. Notably, these phosphorylation events did not enhance MT binding of NuMA FL, suggesting that other phosphorylation sites or kinases may regulate the interaction of NuMA with the MT. It remains to be determined whether NuMA can attain different conformational states through phosphorylation/dephosphorylation and interacting with binding partners for its specific functions.

Our results are largely consistent with an emergent *in vitro* study of the activation of dynein motility and the MT minus-end recognition of NuMA^67^. Unlike our observation that NuMA FL is autoinhibited, Colombo et al. observed more robust MT interaction and dynein activation of NuMA FL^67^. Discrepancies between these studies may be relevant to differences in construct design and clustering of NuMA, or the usage of nonionic detergent to solubilize NuMA FL^67^, which may potentially affect autoinhibitory interactions.

NuMA is a unique dynein adaptor that interacts with MTs at its “cargo binding end”, thereby playing a central role in focusing the minus-ends of MTs^28, 52^. In spindle poles, most MTs are not directly anchored to the centrosomes^2^ but held together by NuMA^4, 9, 52, 68^. Because spindle poles recruit other MT motors, capping proteins, and crosslinkers, it is challenging to dissect the precise role of NuMA and dynein in the minus-end focusing of MTs. We demonstrated that the C-terminal region enables crosslinking of freely floating MTs, presumably due to its clustering and binding to MTs at multiple sites. Neither NuMA nor dynein/dynactin alone could focus or align MTs, but together, they were able to focus minus-ends of MTs in the absence of other proteins. Aster formation requires the accumulation of NuMA at the MT minus-end, consistent with the ability of NuMA to localize to the spindle poles, albeit less efficiently, in the absence of dynein^9^. Transport of these MTs by DDN complexes as cargo towards the minus-end of another MT results in focusing and aligning of MT minus-ends into asters with a large accumulation of NuMA at its center. *In vivo* studies showed that focusing MTs into spindle poles also requires minus-end-directed kinesin-14, KIFC1 (HSET)^52^ and MT-associated crosslinkers^69^. The *in vitro* reconstitution assay we developed is poised to provide molecular insight into the specific contributions of MT motors, crosslinkers, and adaptors to spindle pole formation.

## Extended Data Tables

**Extended Data Table 1.**
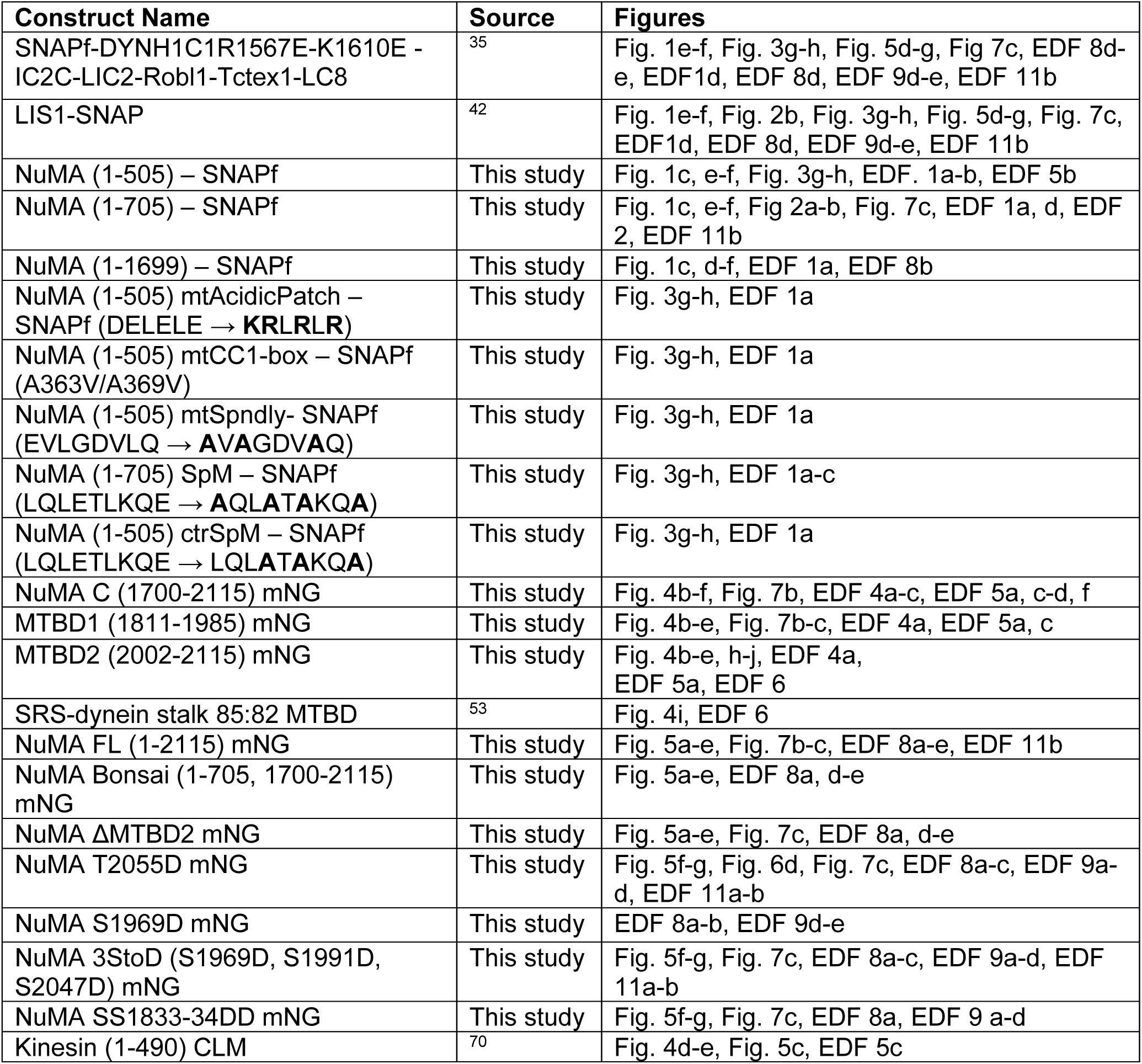
Recombinant protein constructs that were used in this study. (EDF: Extended Data Figure; mNG: mNeonGreen).

**Extended Data Table 2.** Cryo-EM data collection and refinement statistics for the DDN complex.

|  | **DDNL consensus** | **Pointed end with NuMA** |
| --- | --- | --- |
| **EMDB ID** | 52171 | 52172 |
| **PDB ID** | 9HHL | - |
| **Data collection** |  |  |
| Voltage | 300 kV | 300 kV |
| Electron exposure | 40 e<sup>-</sup>/Å<sup>2</sup> | 40 e<sup>-</sup>/Å<sup>2</sup> |
| Magnification | 81000x | 81000x |
| Pixel size | 1.059 Å/px | 1.059 Å/px |
| Defocus range | -1.2 to -2.2 µm | -1.2 to -2.2 µm |
| Total micrographs | 44,287 | 44,287 |
| Initial particle images | 1,418,923 | 1,418,923 |
| Final particle images | 8,092 | 8,092 |
| Map resolution | 6.53 Å | 6.78 Å |
| FSC threshold | 0.143 | 0.143 |
| Sharpening B-factor | -202 | -380 |
| **Model refinement** |  |  |
| Model Resolution | 8.8 | - |
| FSC threshold | 0.5 | - |
| **Model composition** |  | - |
| Non-hydrogen atoms | 64693 | - |
| Protein residues | 13016 | - |
| Ligands | Zn: 3 ADP: 8 ATP: 2 | - |
| **RMSD deviations** |  |  |
| Bond lengths | 0.004 Å | - |
| Bond angles | 1.190 ° | - |
| **Validation** |  |  |
| Clashscore | 3.53 | - |
| MolProbity score | 1.33 | - |
| Rotamers outliers (%) | 0.00 | - |
| **Ramachandran plot** |  |  |
| Favored | 96.87 % | - |
| Allowed | 3.10 % | - |
| Outliers | 0.03 % | - |

**Extended Data Table 3.** Cryo-EM data collection and refinement statistics for the MTs co-decorated with NuMA MTBD2 and dynein-MTBD.

|  | **MT – NuMA MTBD2 – Dynein MTBD** |
| --- | --- |
| EMDB ID | XXX |
| Microscope | Arctica |
| Voltage (kV) | 200 |
| Camera | K3 |
| Defocus range (μm) | -0.8-1.8 |
| Automation software | SerialEM |
| Frames | 50 |
| Total dose (electrons/Å<sup>2</sup>) | 50 |
| Pixel size (Å/pixel) | 1.14 |
| Number of micrographs | 1815 |
| Starting number of particles (pre-symmetry expansion) | 242,128 |
| Number of particles in final map (post-symmetry expansion) | 576,069 |
| Map sharpening B factor (Å) | -40 |
| Map sharpening methods | Relion 5 |
| Symmetry | C1 |
| Overall resolution (Å) | 3.3 |
| Resolution range of map (min-75th percentile) (Å) | 3.1-5.9 |

## Extended Data Figures

**Extended Data Figure 1.**
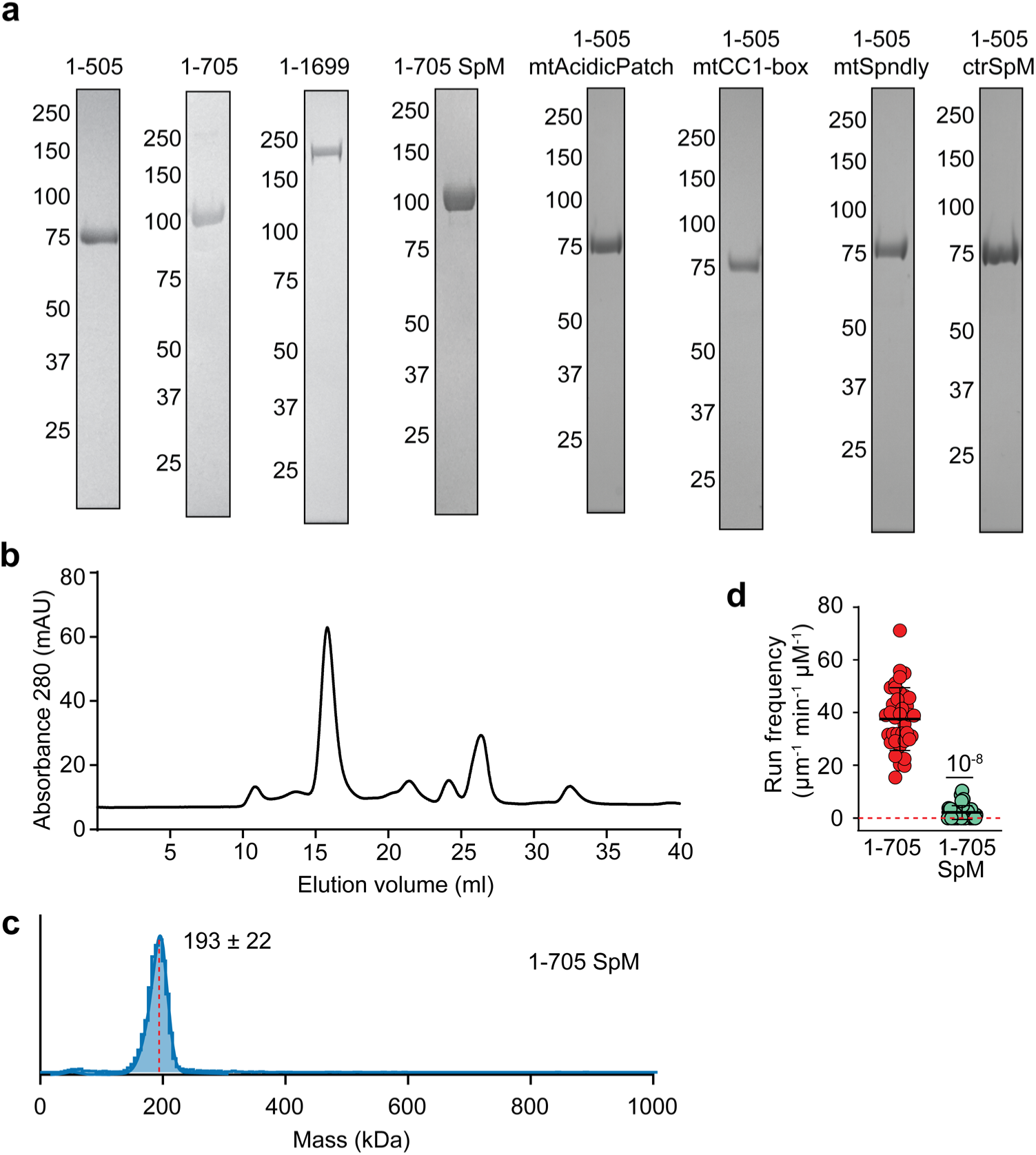
Expression and biochemical characterization of N-terminal NuMA constructs. **a.** Denaturing gel pictures of N-terminal NuMA constructs after gel filtration. The numbers on the left represent molecular weight in kDa. **b.** Elution of NuMA 1-505 from a Superose 6 gel filtration column. **c.** Mass photometry shows that NuMA 1-705 SpM forms a homodimer (mean ± s.d.). **d.** The comparison of the run frequencies of single DDN complexes assembled with NuMA 1-705 and NuMA 1-705 SpM. The centerline and whiskers represent mean and s.d., respectively (from left to right, n = 29 and 40 MTs). The *P* value is calculated from a two-tailed t-test. The velocity and run length of complexes formed with NuMA 1-705 SpM could not be determined due to the infrequency of processive runs.

**Extended Data Figure 2.**
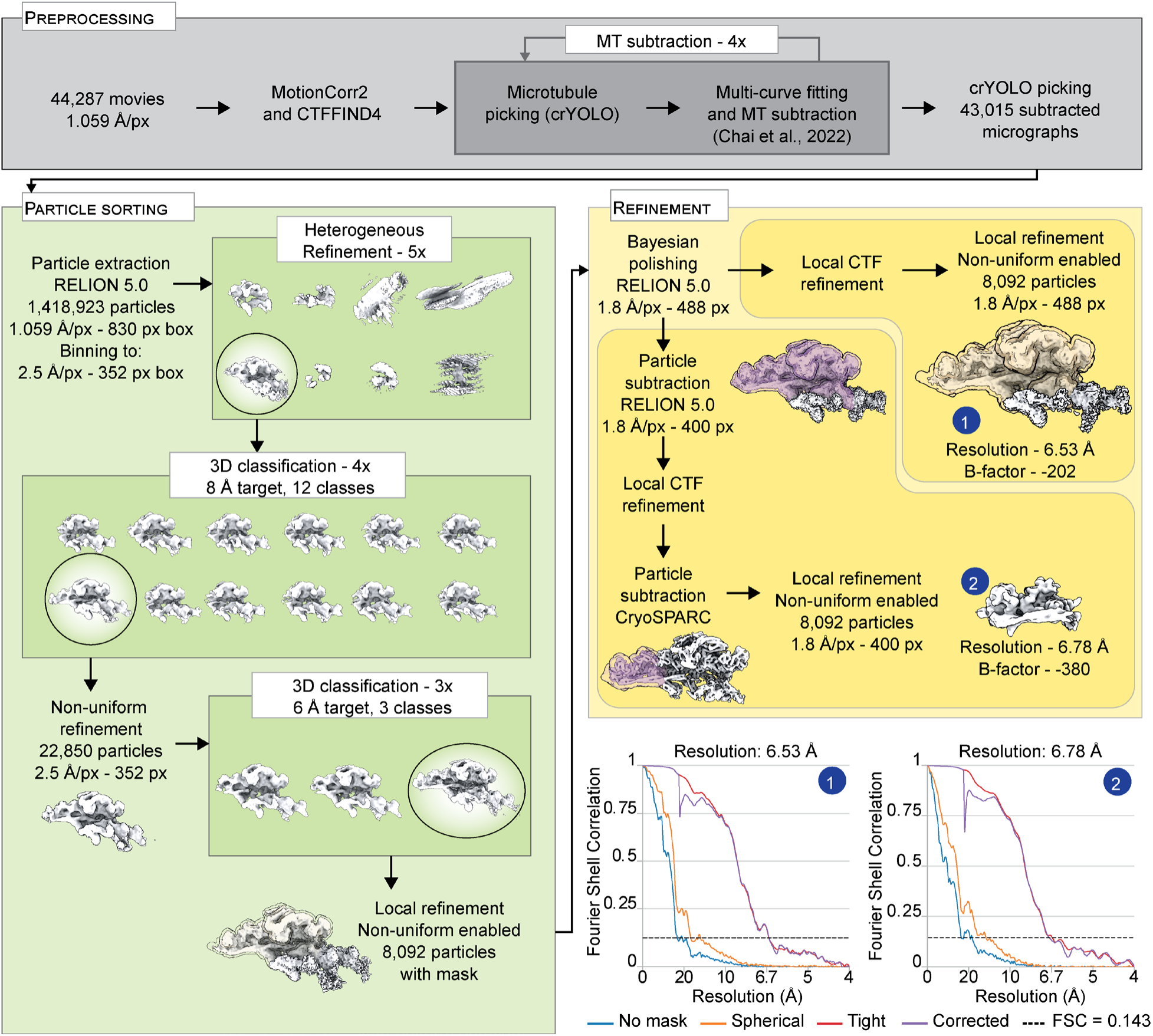
Cryo-EM image processing pipeline for DDN complexes in the presence of Lis1. All refinements and classifications were performed in CryoSPARC unless otherwise specified. The masks used for particle subtraction are displayed in purple. Static masks used for refinements are displayed in yellow. When no mask is displayed, a dynamic mask was used instead. Before local contrast transfer function (CTF) refinement, a non-uniform refinement was performed (not shown). The plots show gold-standard Fourier shell correlation for the final maps.

**Extended Data Figure 3.**
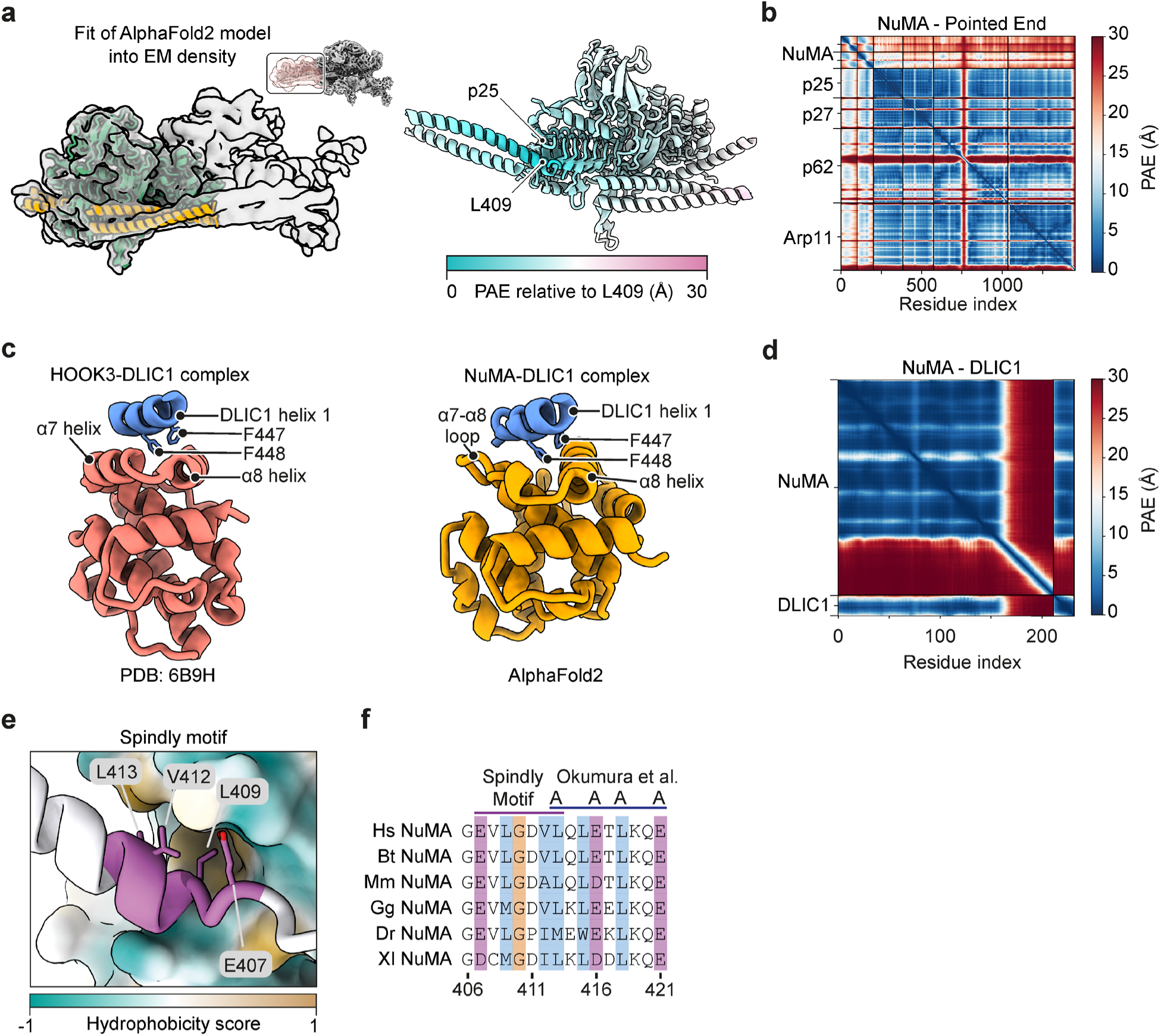
Structural analysis of the interactions between NuMA and dynein-dynactin. **a.** Fitting of an AF2 model of NuMA 351-450 bound to the dynactin pointed end complex into a locally refined EM density for the same region of dynactin, with the mask used and model colored by local predicted alignment error (PAE) values. Residues with low predicted local distance difference test (pLDDT) scores are hidden. **b.** The PAE plot for the NuMA-Pointed end model in a. **c.** Comparison between the AF2 model of the NuMA Hook domain interaction with DLIC1 and the published structure of the complex between the HOOK3 hook domain and DLIC1 (PDB 6B9H)^47^. Residues outside of the HOOK domain and DLIC are not displayed. **d.** The PAE plot for the NuMA-DLIC1 model in c. **e.** The Spindly motif of NuMA bound to the p25 subunit of dynactin, colored by hydrophobicity. **f.** Sequence alignment of the Spindly motif region in NuMA orthologs, with the location of the mutations from Okumura et al.^18^.

**Extended Data Figure 4.**
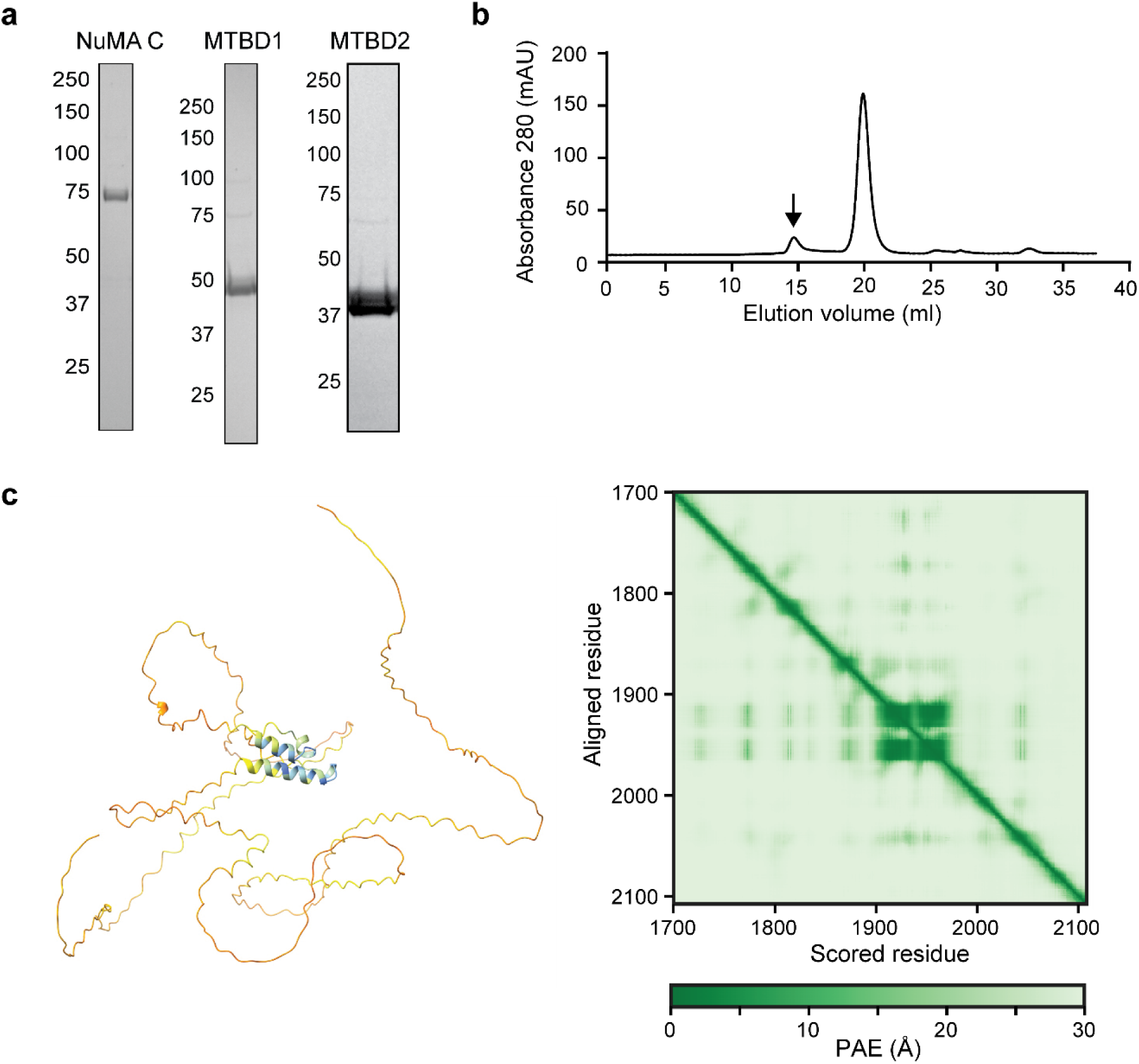
Biochemical characterization and structure prediction of C-terminal NuMA constructs. **a.** Denaturing gel pictures of C-terminal NuMA constructs after gel filtration. **b.** The elution profile of NuMA C (black arrow) from a Superdex 200 gel filtration column. **c.** (Left) AF2 predicts that the NuMA C-terminus is almost fully unstructured except for a single short helix in MTBD1. (Right) PAE for the model.

**Extended Data Figure 5.**
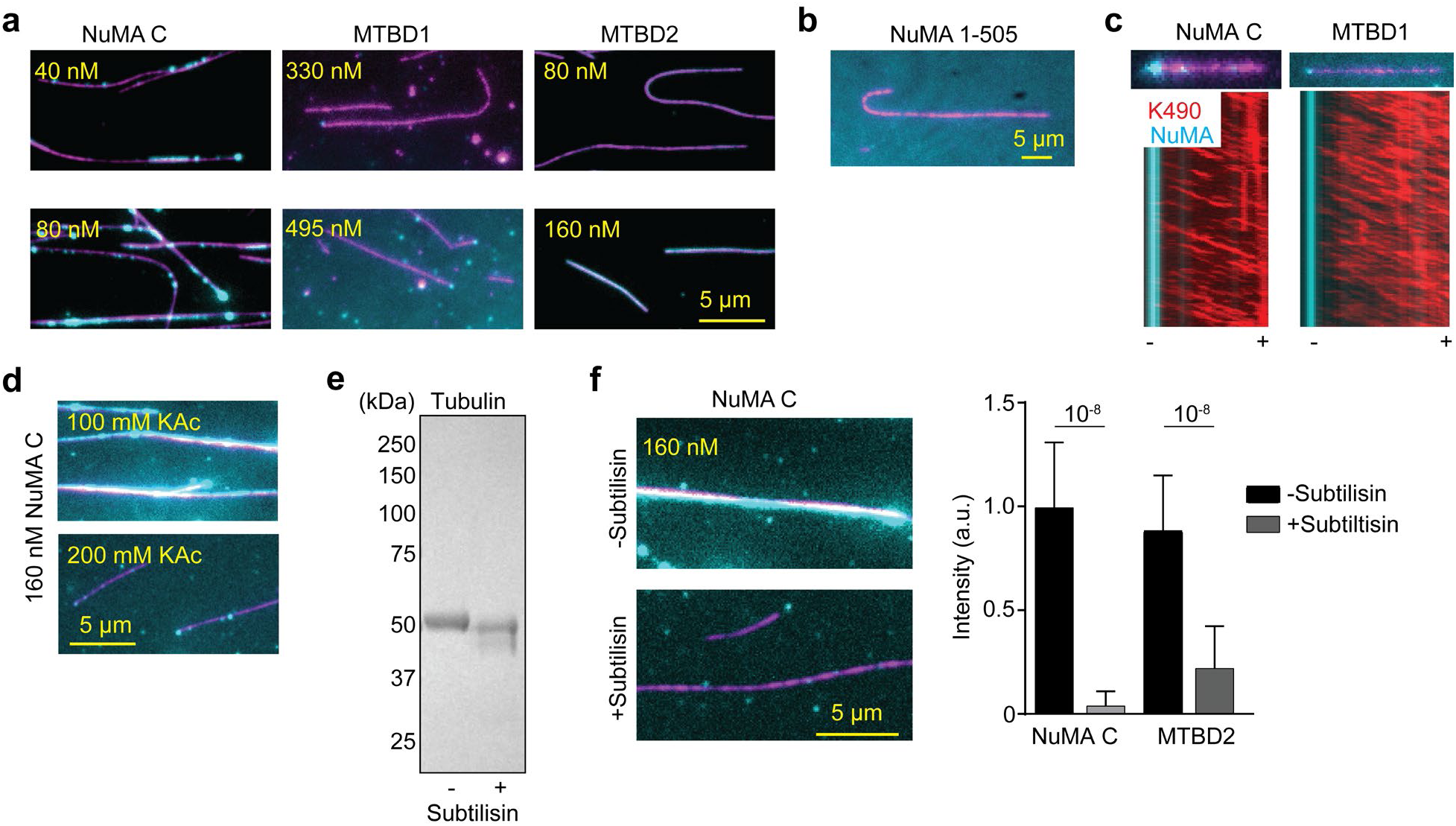
MT binding of C-terminal NuMA constructs. **a.** Example pictures show NuMA C and MTBD2, but not MTBD1, densely decorated surface-immobilized MTs. **b.** An example picture shows that NuMA 1-505 (cyan) in the flow chamber does not bind to MTs (magenta). **c.** Two-color kymographs of tail-truncated kinesin-1 (K490; red) and NuMA C or MTBD1 (cyan) on the MT. MT polarity was determined from the plus-end-directed motility of K490 motors. NuMA accumulates at the minus-end of the MTs. **d.** MT binding of NuMA C is greatly reduced by increasing the salt concentration from 100 mM to 200 mM. **e.** The denaturing gel picture shows the reduction of the molecular weight of tubulin due to the cleavage of tubulin C-terminal tails by subtilisin treatment. **f.** (Left) NuMA C exhibits little to no binding to subtilisin-treated MTs. (Right) The normalized fluorescence intensity of 300 nM NuMA C and 50 nM MTBD2 on surface-immobilized MTs in the presence and absence of subtilisin treatment. Error bars represent s.d. (from left to right, n = 119, 166, 70, and 174 MTs). *P* values are calculated from a two-tailed t-test.

**Extended Data Figure 6.**
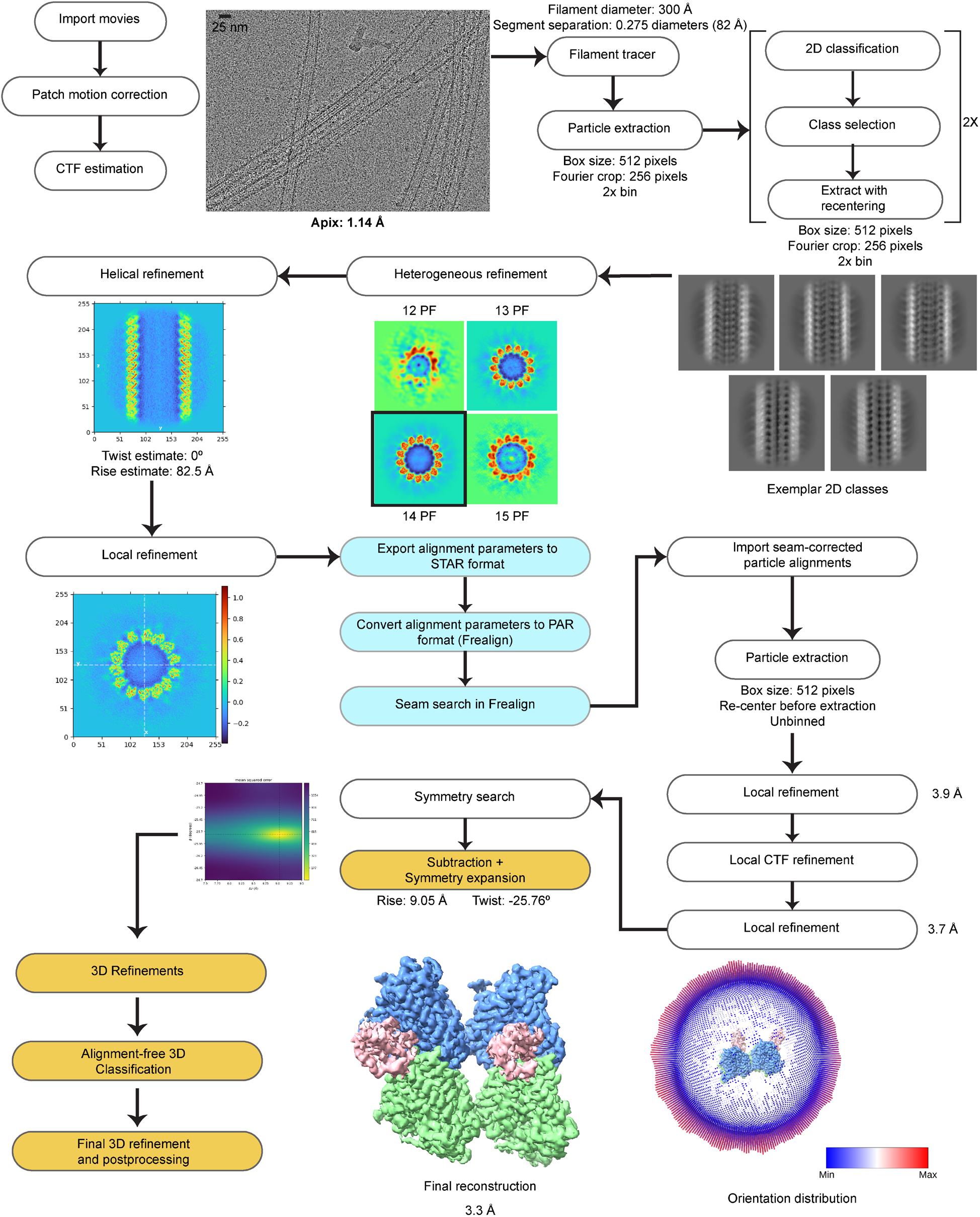
Cryo-EM data processing pipeline used to determine the structure of MTs co-decorated with dynein-MTBD and NuMA MTBD2. Steps with a white background were performed in CryoSPARC; light blue indicates the seam search protocol, and orange highlights steps carried out in RELION 5.

**Extended Data Figure 7.**
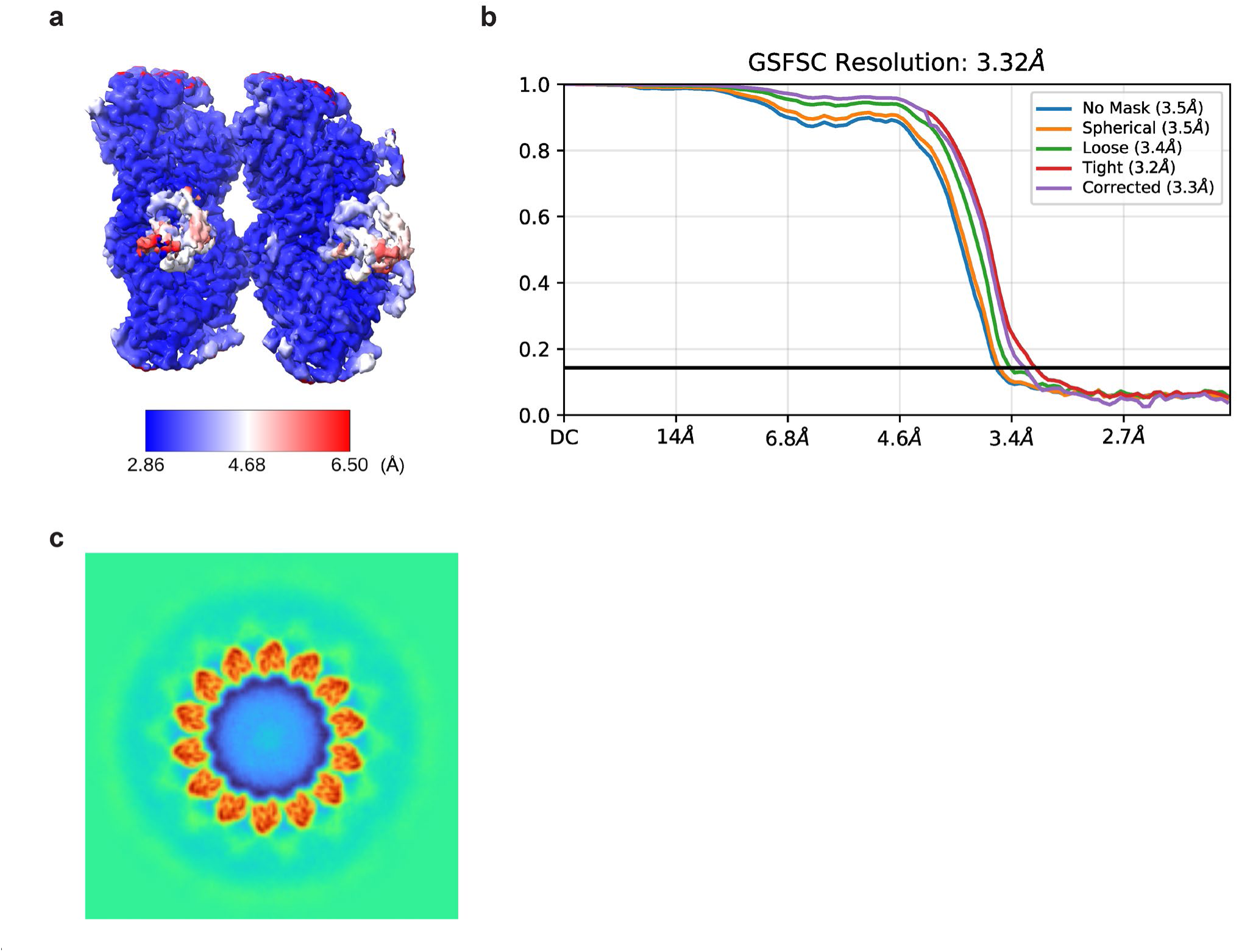
Resolution of the cryo-EM maps. **a.** Local resolution map of the symmetrized reconstruction of MTs co-decorated with dynein-MTBD and NuMA MTBD2. **b.** Fourier shell correlation (FSC) plot of the final map. **c.** An example 3D class of MTs decorated with NuMA MTBD2-mNG in the absence of dynein MTBD. A fuzzy density outside the MT surface corresponds to mNG that is fused to MTBD2, implying that the protein is bound to the MT.

**Extended Data Figure 8.**
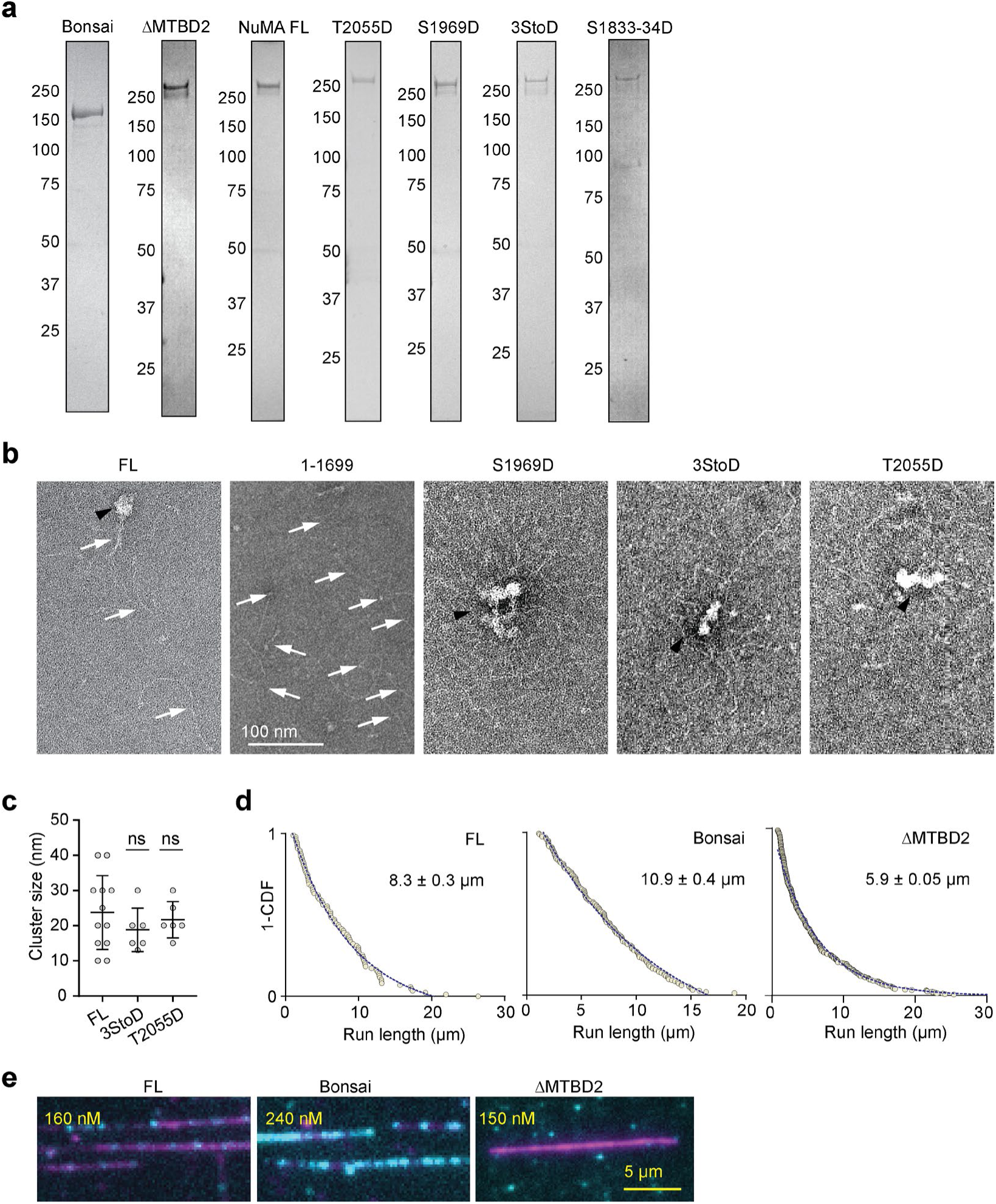
Expression and characterization of NuMA FL constructs. **a.** Denaturing gel pictures of NuMA constructs after gel filtration. **b.** An example negative-stain EM micrograph shows that NuMA FL constructs form large clusters (black arrowheads) with coiled-coils pointing outward (white arrows). In comparison, NuMA 1-1699 forms only elongated coiled coils. **c.** The size of the clusters formed with NuMA constructs in a. The centerline and whiskers represent mean and s.d., respectively (from left to right, n = 12, 6, and 6 clusters). *P* values are calculated from a two-tailed t-test. **d.** The run lengths of single DDN complexes assembled with NuMA FL, Bonsai, and ΔMTBD2 constructs (from left to right, n = 85, 154, and 360 motors). 1-CDFs of motor run length were fitted to a single exponential decay (blue dashed curves) to determine the mean run length (±s.e.). **e.** Example pictures show binding of NuMA FL, Bonsai, and ΔMTBD2 (cyan) to MTs (magenta).

**Extended Data Figure 9.**
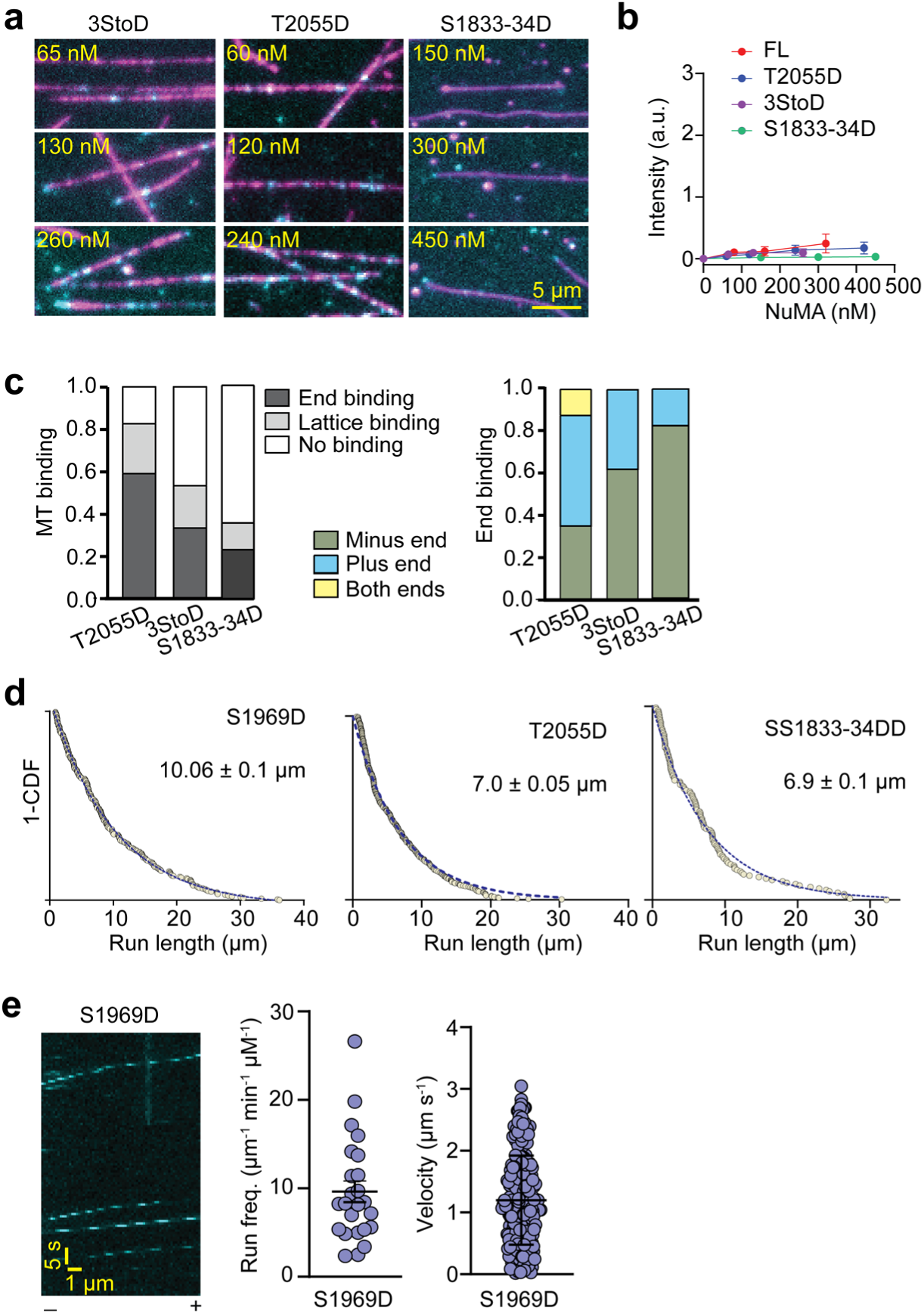
MT binding and DDN motility of phosphomimetic mutants of NuMA. **a.** Example pictures show MT (magenta) binding of NuMA 3StoD and T2055D (cyan) under different concentrations. **b**. The intensity of mitotically phosphorylated NuMA constructs per length of an MT. The centerline and whiskers represent mean and s.d., respectively (from left to right, n = 30, 203, 203, 348 MTs for NuMA FL; 30, 362, 394, 349, and 336 for T2055D; 30, 296, 301, 304 for 3StoD; 25, 156, 92, and 186 for S1833/34D). **c.** Normalized MT binding and end binding preference of mitotically phosphorylated NuMA constructs (from left to right, n = 178, 150, and 272 MTs). **d.** The run lengths of single DDN complexes assembled with mitotically phosphorylated NuMA constructs (from left to right, n= 243, 377, and 136 motors). 1-CDFs of motor run length were fitted to a single exponential decay (blue dashed curves) to determine the mean run length (±s.e.). **e.** An example kymograph, run frequency, and velocity of single DDN complexes assembled with S1969D. The centerline and whiskers represent mean and s.d., respectively (n= 24 MTs for the run frequency and 243 motors for velocity measurements).

**Extended Data Figure 10.**
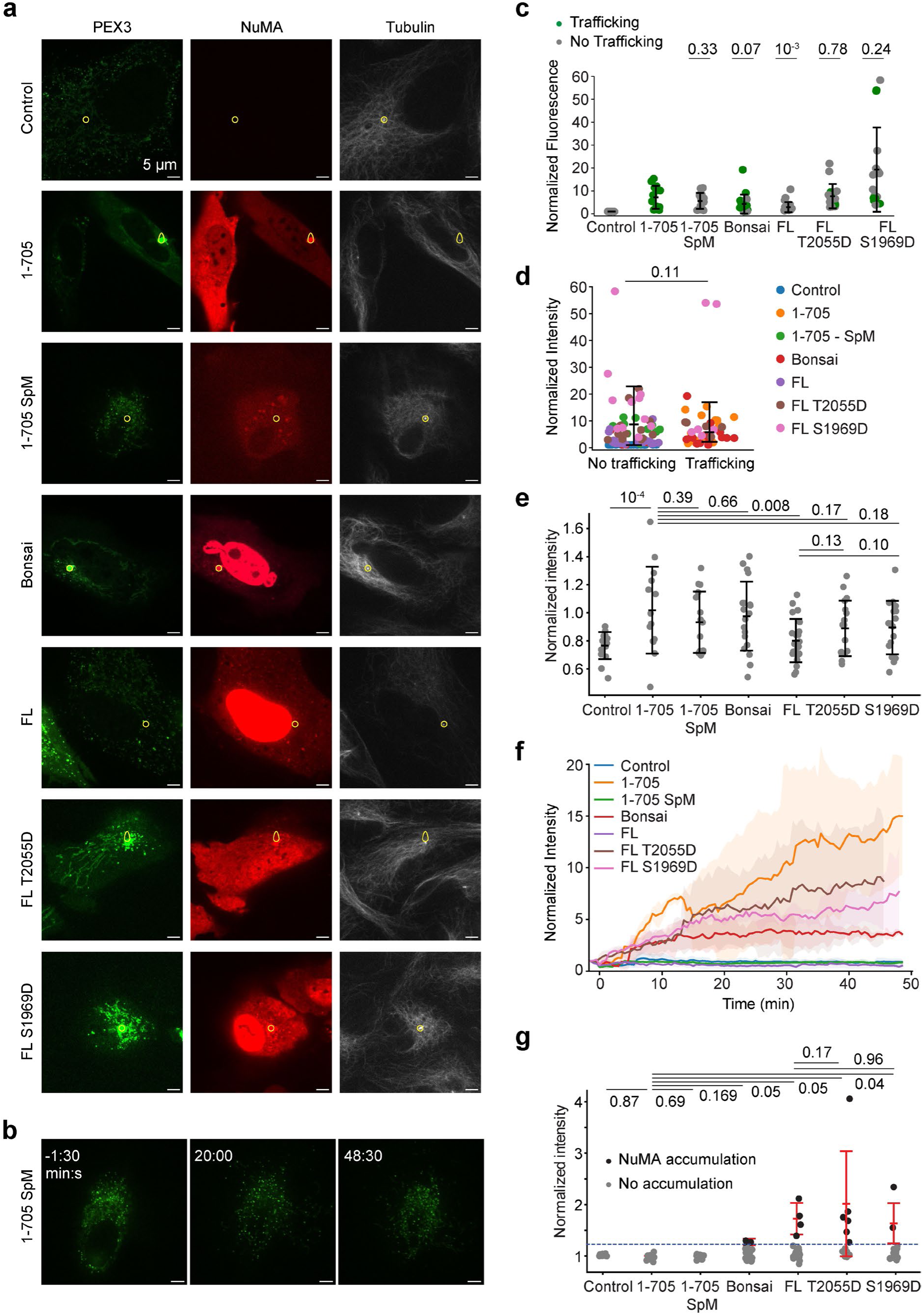
The analysis of peroxisome trafficking assays. **a.** Representative confocal images of peroxisomes (PEX3, green), NuMA (red), and tubulin (SiR-tubulin, grey) from z-stacks taken 45 minutes after rapamycin addition in WT RPE1 cells (control) and RPE1 cells expressing different NuMA constructs. Centrosomes (white dots in the tubulin channel) and consequently the area of quantification is marked by the yellow circle in each still. Cells are the same as in Fig. 6b. **b.** Representative confocal images of peroxisomes (PEX3, green) before and after rapamycin addition in RPE1 cells expressing NuMA 1-705 SpM. Scale bar = 5 µm. **c.** Normalized cytoplasmic NuMA concentration was calculated by averaging the mean pixel intensity for NuMA in the cytoplasm from the timepoints before rapamycin addition. **d.** Normalized cytoplasmic NuMA concentration sorted by trafficking outcome. **e.** The ratio of the final (45 min after rapamycin addition) mEmerald intensity to the initial mEmerald intensity in cytoplasm. No cells showed cytoplasmic accumulation over the threshold (2). **f.** Average peroxisome signal accumulation over time using normalized mEmerald intensity, from a subset of cells where the centrosome (marked by SiR-tubulin) stayed in focus for the majority of frames. For NuMA 1-705, Bonsai, T2055D, and S1969D, only cells that had trafficking were tracked. Time points where the centrosome was out of focus were excluded. Shading represents s.d. From top to bottom, n = 7, 4, 5, 4, 2, 3, and 3 cells pooled from two independent experiments. **g.** Normalized centrosomal NuMA concentration was calculated by averaging the mean pixel intensity for NuMA at the centrosome for the timepoints before rapamycin addition. Cells were scored as either NuMA accumulation (black) if the normalized intensity is above 1.2, or no accumulation (grey) if below the threshold. In c, d, e, and g, n = 16, 14, 15, 19, 22, 17, and 17 cells from left to right (two independent experiments), and the centerline and whiskers represent mean and s.d., respectively. *P* values are calculated from a two-tailed t-test.

**Extended Data Figure 11.**
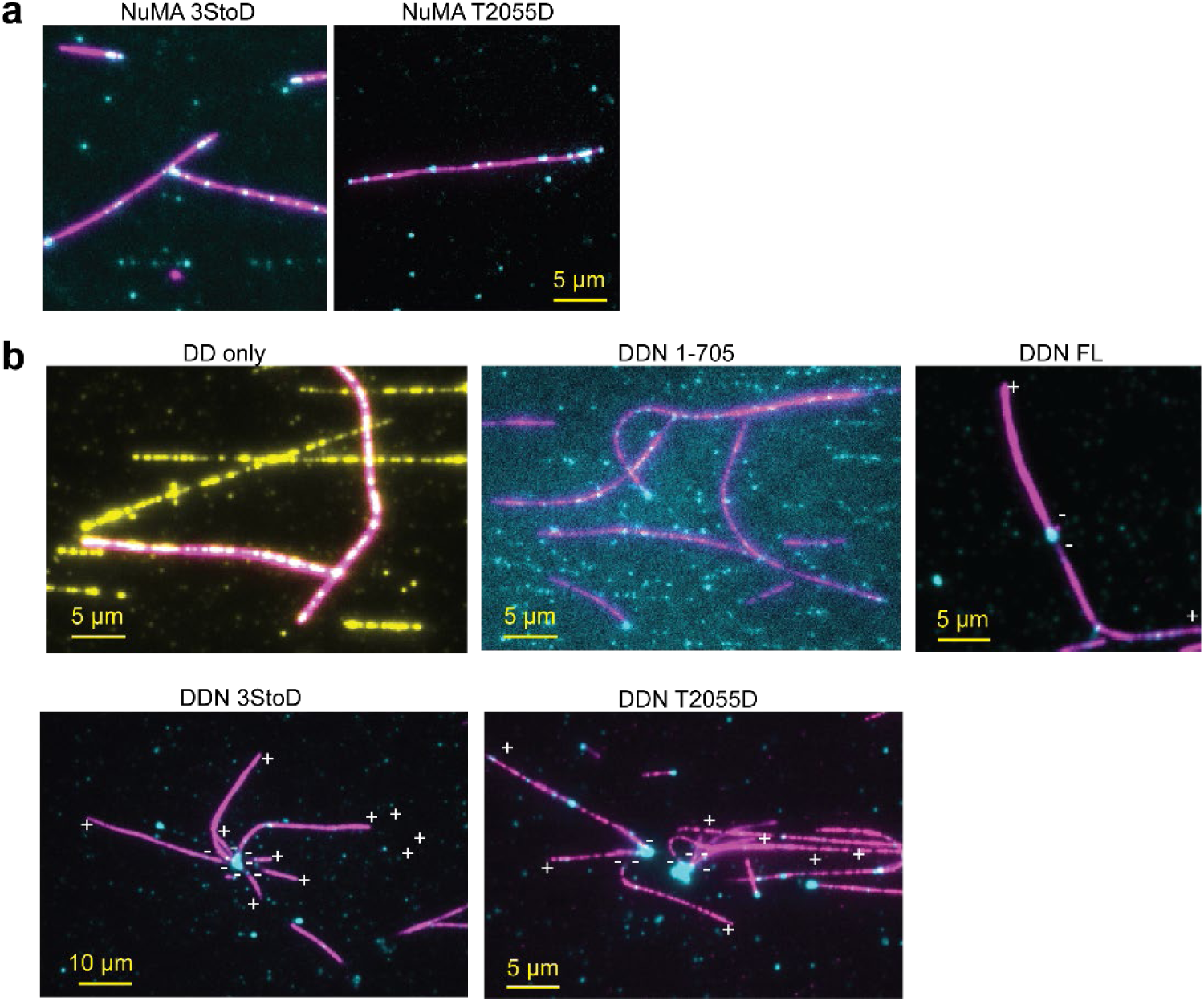
Additional examples of the organization of MTs into aster-like structures by DDN. **a.** NuMA 3StoD and T2055D bind to freely diffusing MTs, but cannot bundle MTs or form MT asters in the absence of dynein/dynactin. **b.** Additional examples of the organization of MTs into aster-like structures by active DDN complexes formed by NuMA 3StoD and T2055D. These structures were not observed in the absence of NuMA (DD only) or the presence of NuMA FL and NuMA 1-705. MT polarity was determined from the plus-end-directed motility of Cy5-labeled K490 (not shown). Dynein, NuMA, and MTs are colored yellow, cyan, and magenta, respectively.

## Materials and Methods

### Cloning and plasmid generation

All human NuMA constructs were cloned into the pOmnibac vector, incorporating an N-terminal His6-ZZ tag followed by a TEV protease cleavage site for protein expression and purification and a C-terminal SNAP or mNG sequence for fluorescent imaging. Full-length and C-terminal NuMA constructs were cloned into a vector containing an additional N-terminal MBP tag between the TEV cleavage and the ZZ tag to improve the solubility of expressed proteins. For live cell imaging, the coding sequence of NuMA-HaloTag-FRB constructs (NuMA 1-705, NuMA-Bonsai, NuMA 1-705 SpM, NuMA-FL, NuMA-FL-T2055D, and NuMA-FL-S1696D) were cloned into a puromycin-resistant lentiviral vector (Addgene #114021). The list of constructs used for protein expression in this study is shown in Extended Data Table 1.

### Cell culture

hTERT-RPE1 cells were purchased from ATCC (CRL-4000). Lentivirus was produced in HEK293T cells that were purchased from ATCC (CRL-3216). All cells were grown in DMEM/F12 media (ThermoFisher) supplemented with 10% tetracycline-screened FBS (PS-FB2; Peak Serum) and maintained at 37 °C and 5% CO_2_.

### Lentiviral plasmids, cell line constructions, and transfections

Lentivirus for each NuMA construct was produced in HEK293T cells. To generate stable polyclonal cell lines, wild-type RPE1 cells were infected with lentivirus and selected using 5 µg mL^-1^ puromycin. PEX3-mEmerald-FKBP (a gift from S. Reck-Peterson, UCSD) was transiently transfected into RPE1 cells to label peroxisomes for 2 or 3 days using Viafect (Promega) according to the manufacturer’s recommendations. Tubulin (including the centrosome) was labeled in cells by adding 100 nM SiR-tubulin and 10 µM verapamil for 60 min before imaging (Cytoskeleton). NuMA was labeled with 100 nM Janelia Fluor 549 (Promega) for 15 min before imaging. Rapamycin was added to a final concentration of 1 µM to induce dimerization of FRB-FKBP.

### Protein expression and purification

For baculovirus production, pOmnibac plasmids were transformed into DH10Bac competent cells and plated on Bacmid plates containing Bluo-Gal for 48 h at 37°C. Selected white colonies were grown overnight at 37°C in 2xYT medium. Bacmid DNA was purified using isopropanol precipitation and transfected into adherent SF9 cells (10⁶ cells mL^-1^) using FuGENE to produce P1 virus. The cells were cultured for 5–6 days at 27°C, and the P1 virus was harvested when more than 90% of the cells were transfected, as determined by monitoring fluorescent gene expression with a fluorescent light microscope. P2 virus was generated by infecting 50 ml of SF9 cells in suspension with 2 ml of P1 virus for 72 h at 27°C. The P2 virus was harvested by collecting the supernatant after centrifugation of SF9 cells at 4,000 g for 5 min at 4°C.

SF9 cells in 1 lt suspension were infected with 1% (v/v) P2 virus and cultured for 72 h at 27°C. The cells were collected by centrifuging at 4,000 g for 10 minutes and washed with ice-cold PBS before being flash-frozen for further purification. Cell pellets were lysed in a lysis buffer supplemented with 1 mM DTT, 2 mM PMSF, and two cOmplete protease inhibitor cocktail tablets (Roche) per liter of cells using a Dounce homogenizer. The lysate was clarified by centrifugation at 65,000 g for 30 minutes, and the supernatant was incubated with IgG Sepharose beads (Cytiva) for 1 hour at 4°C on a rotator. The beads were loaded onto a gravity-flow column and sequentially washed with a lysis buffer, followed by a wash buffer. The beads-protein mixture was then incubated with 0.1 mg/ml TEV protease overnight at 4°C. Proteins were eluted from the column using gravity flow and concentrated with molecular weight cutoff (MWCO) concentrators (Amicon). Protein concentration was determined by measuring absorbance at 280 nm using a NanoDrop 1000 (Thermo Fisher). Proteins containing a SNAP tag were fluorescently labeled by incubating them with a 5-fold molar excess of BG-LD655 or LD555 dye (Lumidyne). Excess dye was removed by gel filtration.

Dynein-expressing cell pellets were lysed in 50 mM HEPES (pH 7.4), 100 mM NaCl, 2 mM MgCl₂, 0.1 mM ATP, and 10% glycerol. The beads-protein mixture was washed with a buffer containing 50 mM Tris-HCl (pH 7.4), 150 mM KAc, 2 mM Mg(Ac)_2_, 1 mM EGTA, 0.1 mM ATP, and 10% glycerol. The protein was then loaded onto a TSKgel G4000SWXL size exclusion column (Tosoh) for further purification.

KIF5B (1-490) was purified using a lysis buffer consisting of 50 mM HEPES (pH 7.4), 1 M NaCl, 1 mM DTT, 2 mM PMSF, 2 mM MgCl₂, 0.1 mM ATP, and 10% glycerol. The wash buffer contained 50 mM HEPES (pH 7.4), 300 mM NaCl, 1 mM DTT, 2 mM MgCl₂, 0.1 mM ATP, and 10% glycerol.

Cell pellets expressing NuMA N-terminal constructs were lysed and washed with 50 mM HEPES (pH 7.4), 300 mM NaCl, 1 mM EGTA, and 10% glycerol. The lysate was loaded onto a Superose 6 Increase 10/300 GL (Cytiva) column for size exclusion chromatography in a buffer containing 50 mM HEPES (pH 7.4), 150 mM NaCl, 1 mM EGTA, and 10% glycerol.

For FL NuMA protein purification, cells were lysed in 50 mM HEPES (pH 7.4), 30 mM NaCl, 2 mM MgCl₂, 1 mM EGTA, and 10% glycerol, supplemented with benzonase nuclease (Sigma). The NaCl concentration was then increased to 300 mM before centrifugation. The FL NuMA protein-beads mixture was washed with 50 mM HEPES (pH 7.4), 300 mM NaCl, 1 mM EGTA, and 10% glycerol and loaded onto a Superose 6 Increase 10/300 GL (Cytiva) column for size exclusion chromatography in 50 mM HEPES (pH 7.4), 150 mM NaCl, 1 mM EGTA, and 10% glycerol.

Cells expressing NuMA C-terminal constructs were lysed in 50 mM HEPES (pH 7.4), 1 M KCl, 1 mM EGTA, and 10% glycerol. The lysate was washed with 50 mM HEPES (pH 7.4), 300 mM KCl, 1 mM EGTA, and 10% glycerol and loaded onto a Superdex 200 Increase 10/300 GL (Cytiva) column for size exclusion chromatography in 50 mM HEPES (pH 7.4), 150 mM KCl, 1 mM EGTA, and 10% glycerol.

Mammalian dynactin was purified as previously described from pig brains using SP Sepharose Fast Flow, MonoQ ion exchange columns (Cytiva), and the TSKgel G4000SWXL size exclusion column (Tosoh)^48^.

### Cryo-EM sample preparation for DDN

To prepare MTs, porcine brain tubulin was diluted in BRB80 buffer (80 mM PIPES pH 6.8, 1 mM MgCl_2_, 1 mM EGTA) to a final concentration of 6 µM. GMPCPP was added to a final concentration of 1 mM and polymerization was induced by incubating at 37 °C for 1 h. MTs were pelleted by centrifugation at 20,000 *g* for 8 minutes and resuspended in BRB80, after which MTs were depolymerized by incubating 30 min on ice. A second polymerization round was performed by adding GMPCPP to a final concentration of 1 mM and incubating the solution at 37 °C for 1 h, followed by another round of centrifugation and resuspension in BRB80. Bradford assay (BioRad) was used to estimate the protein concentration, and MTs were diluted to 0.3 mg mL^-1^ immediately before use.

The DDN 1-705 complex was assembled by mixing the purified dynein, dynactin, Lis1, and NuMA in a 1:2:50:50 molar ratio, respectively, in GF150 buffer (25 mM HEPES pH 7.2, 150 mM KCl, 1 mM MgCl_2_, 1 mM DTT), with dynein at a final concentration of 0.14 µM in 5 µL. The assembly solution was incubated on ice for 30 min and subsequently diluted 2-fold to a final volume of 10 µL in GF150. To bind the complex to MTs, 9 µL of the complex was mixed with 5 µL of diluted MTs and 9 µL of S1 buffer (25 mM HEPES pH 7.2, 5 mM MgSO₄, 1 mM EGTA, 2 mM DTT, 25 mM AMPPNP) and incubated at room temperature for 15 min. MT-bound complexes were separated from unbound protein by pelleting at 20,000 g for 8 min, and then resuspended in 80 µL S2 buffer (25 mM HEPES pH 7.2, 35 mM KCl, 5 mM MgSO₄, 1 mM EGTA, 1 mM DTT, 5.3 mM AMPPNP, 0.01% IGEPAL (Millipore-Sigma)). 4 µL of resuspended solution was applied to freshly glow-discharged Quantifoil R2/2 mesh 300 gold grids (Quantifoil) in a Vitrobot IV (ThermoFisher) at 100% humidity and 22 °C, incubated for 45 s, and blotted for 1.5 s at a blot force of −15 before plunge-freezing into liquid ethane.

### Cryo-EM data acquisition for DDN

The sample was imaged using a FEI Titan Krios (300 kV) equipped with a K3 detector and energy filter (20 eV slit size, Gatan) using automated data collection (ThermoFisher EPU). A total of 44,287 movies was acquired at 81,000X magnification (1.059 Å/pixel), with a 2.32 s exposure time, 40 fractions, and fluence of ∼40 e^-^/Å^2^. A 100 µm objective aperture and a defocus range between −1.2 and −2.2 µm were used during image acquisition.

### Cryo-EM image processing for DDN

Global motion correction and the contrast transfer function (CTF) estimation were performed using MotionCor2 and CTFFIND4 within RELION 4.0 respectively^71–73^. MTs were initially picked using crYOLO 1.9^74^ in single particle mode, utilizing a model trained on manually picked micrographs in filament mode. The coordinates were then resampled at a 4 nm interval using a multi-curve fitting script, resampled coordinates were split into short segments of 10 coordinates each, and MT subtraction was subsequently performed, as previously described^75^.

The resulting “once-subtracted” micrographs were fed back into crYOLO for a second, third, and fourth round of MT picking, multi-curve fitting, filament splitting, and subtraction, all using the same parameters as the first round. After the fourth round, the quadruple-subtracted micrographs were utilized to pick complexes using crYOLO. If no MTs were detected following a given round of subtraction, the last successfully subtracted micrograph was used instead for particle picking.

The picked particles were extracted using RELION 4.0 and imported into CryoSPARC^76^, for particle sorting. Each step of sorting was repeated multiple times to minimize the effects of stochastic variations in sorting results, with the best class of all parallel iterations pooled together. Firstly, particles went through a round of heterogeneous refinement using eight reference volumes, out of which six were contaminants, one was a dynein-dynactin-BICD2 complex with two dynein dimers, and one was a dynein-dynactin-BICD2 complex with one dynein dimer. These volumes were previously determined to be effective in isolating *bona fide* dynein-dynactin-adaptor complexes (unpublished results). The one-dynein class degraded during refinement, so only the two-dynein class was taken forward. Heterogeneous refinement was followed by two rounds of 3D classification in cryoSPARC, with masks spanning the entirety of the complex first, and the more stable component afterwards. In each round, the class with the best density for dynein-dynactin-adaptor was taken forward. The particles coming out of the last classification were locally refined and exported into RELION 5.0 for particle polishing and rebinning, after which they were imported again into CryoSPARC for CTF refinement and particle subtraction. A graphical summary of the processing pipeline including particle numbers, FSCs, B-factors, intermediate maps, and masks used for refinements, classifications, and particle subtractions are shown in Extended Data Figure 2.

### Model building and refinement of DDN

Model building was performed using ISOLDE^77^ and COOT^78^, and refinements were performed in PHENIX^79^. Protein Data Bank (PDB) models or AF2 predictions were docked into cryo-EM maps using UCSF ChimeraX^80^. Refinement statistics are available in Extended Data Table 2. AF2 models were generated using a local installation of ColabFold 5^81^, using MMSeq^82^ for homology searches and AlphaFold2-Multimer^83^.

To build the full complex model, we docked coiled-coil fragments from an AF2 model of NuMA (1-705) dimer into the adaptor density based on their length and fitted using restrained flexible fitting in ISOLDE. We could not model the linkers between coiled-coil fragments. We have assigned fragments together based on distance, but we could not exclude that the other possible arrangement might be accurate. The model of dynactin and the dynein tails was built using PDB ID 8PTK^46^ as the starting model. To model the CC3 fragment, including the Spindly motif and the CC1b-CC2 linker, we used an AF2 model of NuMA (351-450) dimer and the pointed end subunits, which was joined to the CC1b and flexibly fitted with restraints into the density. Using the density from the pointed end subcomplex local refinement, we could assign the CC2 making the Spindly box contacts and the CC1b contacting the side of the pointed end to the same molecule. The linker connecting the other protomer was not resolved and deleted at this stage.

### Negative stain EM

Proteins for negative stain were diluted to 20 nM with the following buffer (50 mM HEPES pH 7.4, 150 mM KCl) and 4 µL of sample was applied to homemade carbon-coated grids after glow discharging the grids. Samples were then stained with 1.5% uranyl formate, blotted, and then air dried. Imaging was performed at ∼30,000x magnification with a Tecnai 12 microscope.

### Cryo-EM sample preparation for MTs

Lyophilized porcine brain tubulin (Cytoskeleton) was resuspended to 10 mg/mL-1 in BRB80 buffer supplemented with 10% (v/v) glycerol, 1 mM GTP, and 1 mM DTT. 5 μL of the tubulin solution was polymerized at 37 °C for 15 min. 1 μL of 2 mM taxol was added to the tubulin solution and incubated at 37 °C for 10 min. 1 μL of 2 mM taxol was added again, and the solution was incubated for 30 min. MTs were pelleted by centrifugation at 37 °C and 17,000 rcf for 20 min. The supernatant was discarded, and the pelleted MTs were resuspended in resuspension buffer (BRB80 buffer supplemented with 250 μM taxol). Before sample preparation, all MT-binding proteins were desalted to BRB80 using Zeba Spin columns (Pierce).

To prepare MT-bound SRS-Dynein and NuMA MTBD2 samples, 2.5 μL of 2.5 μM taxol-MTs were incubated on a glow-discharged holey carbon cryo-EM grid (QuantiFoil, Cu 300 R 1.2/1.3) for 30 s, manually blotted with Whatman filter paper. Then, 2.5 μL of 5 μM SRS-Dynein were added and incubated on the grid for 30 s and blotted with Whatman filter paper. Finally, 2.5 ul of 5uM NuMA MTBD2 was added to the grid. The grid was transferred to a Vitrobot (ThermoFisher) set at 22 °C and 100% humidity, plunge-frozen in liquid ethane with a blot force of 5 pN and a blot time of 6 s, and transferred to liquid nitrogen.

### Cryo-EM data collection of MTs

Cryo-EM data were collected on an Arctica microscope (ThermoFisher), operated at 200 kV with a K3 direct electron detector (Gatan). Images were acquired at 36,000x nominal magnification at 1.14 Å/pixel. All data was acquired in the super-resolution mode with a dose rate of ∼7.2 electrons/pixel/second and exposure time of ∼9 s dose-fractionated into 50 frames. All data were collected using the SerialEM software package^84^.

### Cryo-EM image processing for MTs

For the NuMA–dynein dataset, movie stacks were imported into CryoSparc^76^ and subjected to patch-based motion correction. CTF parameters were estimated using the patch CTF job and manually curated to remove poor-quality micrographs. Filaments were auto-picked using the filament tracer, using microtubule 2D class templates. A segment separation of 82 Å was used, corresponding to the length of one αβ-tubulin dimer. Particle images were extracted with a 512-pixel box size and Fourier-cropped to 256 pixels for early processing steps.

Two rounds of 2D classification were performed, and only classes showing clear microtubule features were retained; particles with blurry signal or off-centered filaments were discarded. Microtubules with different protofilament (PF) numbers were separated through a heterogeneous refinement using low-pass-filtered templates corresponding to 12-, 13-, 14-, and 15-protofilament microtubules. The 14-PF population represented the majority and was selected for further analysis.

These 14-PF particles were refined using helical refinement with an initial rise of 82.5 Å and a twist of 0°. Local refinement using a cylindrical mask further improved angular assignments. Seam positions were then determined on a per-particle basis using a Frealign-based routine^85^. CryoSparc alignment files from the last local refinement were converted to STAR format using csparc2star (PyEM, 10.5281/zenodo.3576630) and then to PAR format via a custom Python script. Following seam assignment, particles were re-imported into CryoSparc with updated alignments and re-extracted at full resolution (512-pixel box size), using the refined coordinates to recenter the extraction. A local refinement was performed on this particle set, followed by per-particle local CTF refinement. Symmetry parameters (rise and twist) were estimated using CryoSparc’s symmetry search tool.

The resulting particle stack was then imported into Relion 5 for further processing^86, 87^. Signal subtraction was applied to retain only the density surrounding two tubulin dimers and associated proteins, and this subtracted dataset was symmetry-expanded using the previously determined symmetry parameters. These particles were refined using the same small mask as before, and subjected to alignment-free 3D classification to clean the dataset and resolve compositional heterogeneity. A final consensus reconstruction was obtained following refinement and postprocessing in Relion 5. Cryo-EM processing parameters are provided in Extended Data Table 3.

### Fluorescence microscopy

For *in vitro* imaging, a custom-built multicolor objective-type TIRF microscope was used, equipped with a Nikon inverted Ti-E microscope body, a 100X magnification, 1.49 N.A. apochromatic oil-immersion objective (Nikon), and a Perfect Focus System. Fluorescent signals were detected using an electron-multiplied charge-coupled device (EMCCD) camera (Andor, iXon EM+, 512 × 512 pixels), resulting in an effective pixel size of 160 nm after magnification. The mNG, LD555, and LD655 probes were excited by 488 nm, 561 nm, and 633 nm laser beams (Coherent), respectively. The fluorescence signal was filtered using a notch dichroic filter and 525/40, 585/40, and 695/75 bandpass emission filters (Semrock) for the respective probes. Micro-Manager 1.4 was used to control the microscope and acquire movies.

For live cell imaging, cells were plated onto #1.5 glass-bottom 35 mm dishes coated with poly-D-lysine (P35G-1.5-20-C, MatTek Life Sciences) and imaged in a humidified stage-top incubator maintained at 37 °C and 5% CO_2_ (Tokai Hit). Cells were imaged on a spinning disk (CSU-X1, Yokogawa) confocal inverted microscope (Eclipse Ti-E, Nikon Instruments) with the following components: Di01-T405/488/568/647 head dichroic (Semrock); 488 nm (120 mW), 561 nm (100 mW), and 639 nm (160 mW) diode lasers; ET525/36M, ET600/50M, and ET676/37M emission filters (Chroma Technology); and an iXon3 Ultra 897 EMCCD camera (Andor Technology). Images were acquired with a 100x 1.45 Ph3 oil objective (Nikon Instruments) using Micromanager 2.0.0.

### Flow chamber preparation

For *in vitro* fluorescence imaging assays, polyethylene glycol (PEG)-coated coverslips were used to prevent nonspecific surface binding of proteins. The coverslips were washed with water, acetone, and water again, each for 10 min in an ultrasonic cleaner. For hydroxylation, they were incubated in 1 M KOH for 40 min in the ultrasonic cleaner, followed by rinsing with water and methanol. The coverslips were then treated with an aminosilanization mixture containing 3-aminopropyltriethoxysilane, acetate, and methanol for 10 min, rinsed with water for 1 min, and subjected to a second 10 min incubation in the aminosilanization mixture in the ultrasonic cleaner. The coverslips were then rinsed with methanol and air-dried. Hydroxylated coverslips were biotinylated by placing 30 μl of 25% biotin-PEG-succinimidyl valerate in NaHCO₃ buffer (pH 7.4) between two coverslip pieces, forming a sandwich, and incubating overnight at 4°C. The coverslips were then rinsed with water, air-dried, vacuum-sealed, and stored at −20°C. Flow chambers were prepared by attaching the PEG-biotin-coated coverslip to a glass slide using double-sided tape.

### MT polymerization assays

Biotinylated MTs were prepared by diluting 4 μl of 38 mg/mL unlabeled pig brain tubulin and 2 mg/ml biotin-labeled pig brain tubulin in 56 μl BRB80 buffer (80 mM PIPES, pH 6.8, 1 mM MgCl₂, 1 mM EGTA). This mixture was combined with 60 μl of polymerization buffer (1x BRB80, 2 mM GTP, 20% DMSO) and incubated at 37°C for 45 min. After adding 1 mM taxol, the tubulin mixture was incubated for an additional 45 min at 37°C. The polymerized MTs were pelleted by centrifugation at 21,000 g for 15 min at 37°C and resuspended in 25 μl of BRB80 supplemented with 10 μM taxol and 1 mM DTT. The same procedure was followed to prepare biotin-free, fluorescently labeled MTs by diluting 4 μl of 40 mg/mL pig brain tubulin, 5% of which was Cy3- or Cy5-labeled. For subtilisin treatment, 2 mg/mL polymerized MTs were incubated with 0.025 mg/mL subtilisin for 1 h at 37° C. The proteolytic activity of subtilisin was quenched by adding 1 mM PMSF. The cleavage of the tubulin tails by subtilisin was confirmed using denaturing gel analysis.

For GMPCPP-MT seed preparation, a mixture of unlabeled tubulin, 5% biotinylated tubulin, and 5% Cy3-labeled tubulin was diluted to 3 mg/mL in ice-cold BRB80 containing 10% DMSO. After 3 min incubation on ice, the mixture was centrifuged at 400,000 g for 10 min at 4°C. The supernatant was mixed with GMPCPP to a final concentration of 1 mM and incubated at 37°C for 20 min. The GMPCPP-stabilized MT seeds were pelleted by centrifugation at 400,000 g for 10 min at 37°C. The pellet was gently resuspended in 25 μL of BRB80 at room temperature.

### Single-molecule motility assays

To immobilize biotinylated MTs, the PEG/PEG-biotin coated flow chambers were incubated with 1 mg/ml streptavidin for 3 min, followed by a wash with MB buffer (30 mM HEPES pH 7.4, 5 mM MgSO_4_, 1 mM EGTA, 1 mM DTT, and 10 µM Taxol). The biotinylated MTs were then flowed into the chamber, incubated for another 3 min, and washed with MB buffer. For dynein motility assays, dynein and dynactin were mixed and incubated on ice for 3 min. NuMA was then added and incubated for an additional 3 min, followed by the addition of Lis1 with a further 3 min incubation. The molar ratio of the DDNL components in the stock mixture was 1:3:20:30 (dynein:dynactin:NuMA:Lis1) for N terminal NuMA constructs and 1:3:5:40 for NuMA FL constructs. The final volume of the DDNL stock mixture was adjusted to 10 µL by adding MBC buffer (MB buffer supplemented with 1 mg/mL casein, 0.5% pluronic acid, 100 mM KAc, and 0.4% methylcellulose). The DDNL stock was diluted 10-fold for the FL NuMA construct and 60-fold for the N-terminal NuMA construct. Both dilutions were prepared in imaging buffer (MBC buffer supplemented with 0.1 mg/mL glucose oxidase, 0.2 mg/mL catalase, 0.8% D-glucose, and 1 mM ATP) and introduced into the flow chamber. For DDN motility assays with N-terminal NuMA, 20 nM NuMA and 30 nM Lis1 were used in the flow chamber. For assays with FL NuMA, 20 nM NuMA and 160 nM Lis1 were used. Motility was recorded for 200 frames with 0.2 s exposure per frame.

For aster formation assays, biotinylated MTs were immobilized to the PEG/PEG-biotin surface. The DDNL mix with Cy3-labeled, biotin-free MTs was then flowed into the chamber. The aster formation was imaged by overlaying the mNG-labeled NuMA and Cy3-labeled MT channels. To assess the directionality of the MTs, Cy5-labeled KIF5B (a.a. 1-490) was added to the channel and imaged.

### NuMA MT binding assays

To test the MT binding affinity of NuMA, taxol-stabilized MTs were labeled with Cy3 and biotin and immobilized on PEG/PEG-bio coated coverslips using streptavidin-biotin linkage. Serial dilutions of each mNG fused NuMA construct were flowed into the channel in MBC buffer supplemented with 0.1 mg/mL glucose oxidase, 0.2 mg/mL catalase, and 0.8% D-glucose. Images were taken for both the NuMA and MT channels under each condition.

To test the MT binding preference of NuMA, Cy3-labeled free-floating MTs were mixed with NuMA for 3 minutes, diluted in MBC buffer supplemented with 0.1 mg/mL glucose oxidase, 0.2 mg mL^-1^ catalase, 0.8% D-glucose, and 1mM ATP, and then flowed into a channel. 15 nM Cy5 labeled human kinesin-1 K490 was included in the imaging buffer to determine the directionality of the MTs. Final concentrations of 25 nM NuMA C, 165 nM MTBD1, 4 nM MTBD2, 25 nM NuMA-FL, 25 nM 3StoD, and 25 nM T2055D were used.

### Dynamic MT assays

Biotinylated GMPCPP-MT seeds were immobilized in a 1 mg/mL streptavidin-coated flow chamber by incubating for 3 min. To generate dynamically growing and shrinking MTs, a dynamic MT mixture (2 mg/mL tubulin with 5% Cy3 labeling, 4 mM GTP, 1 mM ATP, 1 mg/mL casein, 0.5% pluronic acid, 1 mM DTT, and 0.2% methylcellulose in BRB80, pH 6.8,) was introduced into the chamber. To assess its effect, 100 nM of the NuMA C-mNG was added to the mixture. MT dynamics was monitored by timelapse imaging with 200 frames, a 0.2 s exposure time, and a 5000 s interval between the frames.

### Mass photometry

High-precision coverslips (Azer Scientific) were cleaned sequentially with isopropanol and water three times, using an ultrasonic cleaner for 2 min per cycle, and then air-dried. Gaskets were rinsed alternately with water and isopropanol three times, air-dried, and placed on a clean coverslip. For autofocus, 14 µL of mass photometry buffer (50 mM HEPES, pH 7.4, 150 mM NaCl, 1 mM EGTA, and 10% glycerol) was loaded into a well of the gasket. The protein sample was diluted to 5–20 nM in mass photometry buffer and added to the autofocused gasket well. Protein contrast counts were measured using a TwoMP mass photometer (Refeyn 2) with a minimum of two replicates. The instrument was calibrated for mass measurements using a mixture of conalbumin, aldolase, and thyroglobulin. DiscoverMP software (Refeyn) was used to fit the mass photometry profiles to multiple skewed Gaussian peaks and to determine their mean, standard deviation, and proportions.

### Peroxisome trafficking assays

Cells expressing both PEX3-mEmerald-FKBP and a NuMA-HaloTag-FRB construct were imaged every 30 s for approximately 50 min. Rapamycin was added to the dishes after 90 s of imaging. At the end of the time-lapse imaging, a z-stack was taken to determine whether peroxisomes had clustered at the centrosome. Peroxisome trafficking was determined by quantifying the ratio of final (45 min) to initial (0 min) PEX3 intensity integrated over 2.54 µm x 2.54 µm oval section at the centrosome using FIJI. Cells were considered to have trafficking if the ratio was above 2. As a control, the final to initial PEX3 intensity in the cytoplasm was also calculated. NuMA intensity in the cytoplasm was calculated over a 10.07 µm x 10.07 µm rectangular area using FIJI, averaged over three timepoints before rapamycin addition, and normalized to control. NuMA intensity at the centrosome was quantified over a 2.54 µm x 2.54 µm oval selection before rapamycin addition, averaged over three timepoints before rapamycin addition, and normalized to the control. Cells were considered to have NuMA accumulation if the normalized centrosomal intensity was above 1.2. In time-lapses where the centrosome remained in focus for the majority of imaging, PEX intensity was integrated over a 2.54 µm x 2.54 µm oval selection at the centrosome at every time point and normalized to the first time point of imaging before rapamycin addition. Individual timepoints where the cell temporarily was out of focus were excluded from the traces.

### Data analysis

Recorded movies of DDN single-molecule motility assays were analyzed using ImageJ. For two-color imaging, the fluorescence channels were merged in ImageJ to create a composite image. A custom ImageJ macro was used to generate kymographs by plotting segmented lines along MTs. For run frequency analysis, the number of processive NuMA molecules on each MT was divided by the MT length (µm), time (min), and NuMA concentration (µM). Dynein’s velocity and run length were determined by identifying the start and end points of each processive NuMA run. For MT binding assays, the intensity of NuMA on the MTs was quantified after background subtraction using a custom MatLab code (MTIMBS). The dynamic MT growth velocity was calculated by drawing kymographs and identifying the start and end points of each growing end in ImageJ.

To determine the binding preference of different NuMA constructs on biotin-free MTs, NuMA fused to mNG, Cy3-labeled MTs, and Cy5-labeled KIF5B (1-490) channels were overlaid. MTs with mNG signal accumulated at one or both ends were manually counted as exhibiting end-binding. The directionality of the MTs was confirmed by analyzing the movement of KIF5B (1-490) through kymographs. NuMA-bound MTs without signal accumulation at the tips were quantified as displaying lattice binding.

### Statistical analysis

*P* values were determined using statistical tests in Prism. Fisher’s exact tests were performed to compare categorical datasets. Two-sided two-sample t-tests were performed to compare continuous datasets, assuming that NuMA and dynein levels are approximately normally distributed. Each result was based on a minimum of two independent experiments. The number of replicates and data points (n) and the specific statistical methods used are provided in the figure legends. Representative data from independently repeated experiments are shown.

### Data Availability

A reporting summary for this article is available as the Supplementary Information file. The main data supporting the findings of this study are available within the article and its Extended Data Figures. Protocols that support the findings of this study can be found in Methods. The constructs that express wild-type and mutant versions of NuMA will be deposited to AddGene. Raw microscopy data will be made available by the corresponding authors upon request. Cryo-EM structures are deposited with PDB and EMDB under accession codes 9HHL/52171 (DDNL consensus) and 52172 (Pointed end with NuMA).

### Code Availability

The custom code used to analyze experimental data is uploaded to the Yildiz Lab code repository (www.yildizlab.org/code_repository) and GitHub (https://github.com/Yildiz-Lab/MTIMBS).

## Supplementary Movie Legends

**Supplementary Movie 1. The NuMA N terminus activates dynein motility.** One color TIRF imaging of representative DDN N-terminal complexes on surface-immobilized unlabeled MTs in the presence of Lis1. NuMA was labeled with LD555, and the images were acquired at 200 ms exposure time per frame.

**Supplementary Movie 2. FL NuMA is autoinhibited and phosphorylation through its C-terminus activates DDN motility.** Single color TIRF imaging of DDN FL complexes on surface-immobilized unlabeled MTs in the presence of Lis1. The motility was monitored by the fluorescence signal of NuMA-mNG at 200 ms exposure time per frame.

**Supplementary Movie 3. NuMA can activate dynein in interphase cells.** Timelapse confocal imaging of a representative WT RPE1 control cell, transfected with PEX3-mEmerald-FKBP (grey), infected or transfected with various NuMA-HaloTag-FRB constructs (not shown), and stained with SiR-tubulin (not shown), with rapamycin addition at time 0:00 (min:s). Peroxisomes randomly diffuse throughout the cytoplasm in control and in conditions where the NuMA construct does not activate dynein. Peroxisomes traffic to the centrosome, as defined by SiR-tubulin staining, when the NuMA construct activates dynein. PEX3-mEmerald-FKBP levels vary between cells due to variations in transfections and imaging of individual cells on different days post-transfection. The movie corresponds to still images in Figure 6b. Scale bar = 5 µm.

**Supplementary Movie 4. Active NuMA FL forms asters by focusing the minus-ends of MTs with dynein/dynactin.** Two-color imaging of DDN 3StoD motility on surface-immobilized unlabeled MTs in the presence of free-floating Cy3-labeled MTs (magenta) and Lis1. The motility was monitored by the fluorescence signal of NuMA-mNG (cyan). The images were acquired at 200 ms exposure time per frame.

## Acknowledgments

We are grateful to the members of the Yildiz laboratory for helpful discussions and carefully reading the manuscript, E. Nogales for negative stain EM imaging, S. Reck-Peterson for reagents, the MRC-LMB EM Facility for access to and support for EM sample preparation and data collection, and J. Grimmett, T. Darling, and I. Clayson of MRC-LMB Scientific Computing for scientific computing resources, CalCryo facility, D. Toso, and P. Tobias for cryoEM sample preparation and data collection. This work was funded by grants from the NIH (GM136414), and NSF (MCB-1055017, MCB-1617028) to A.Y., NIH F31CA275394, AHA Predoctoral fellowship (908941) and a UCSF Discovery Fellowship to N.H.C., grants from the NIH (R35GM136420) and NSF (1548297) to S.D., NSF Graduate Research Fellowship (DGE-2146752) to A.T., Wellcome Trust (227434/Z/23/Z and 210711/Z/18/Z), MRC (MC_UP_A025_1011) to A.P.C, and UK Research and Innovation Horizon Europe Funding Guarantee (101105484) to E.A.d’A. S.D. is a Chan Zuckerberg Biohub Investigator.

## Author Contributions Statement

M.A., A.Y., A.P.C., and S.D. conceived the study and designed the experiments. M.A. prepared the constructs, and isolated and labeled the proteins. M.A. and X.Z. performed functional assays in vitro. M.A. and Y.Z. measured MT dynamics. N.H.C. and M.B. performed live cell imaging assays. E.A.d’A. performed cryo-EM imaging of the DDN complex. A.T. performed negative-stain EM experiments. A.T. and J.M.P.B. performed cryo-EM imaging of NuMA decorated MTs. M.A., E.A.d’A., N.H.C., A.T., and J.M.P.B. performed image processing and data analysis. M.A., A.Y., E.A.d’A., N.H.C., S.D., and A.P.C. wrote the manuscript, and all authors read and commented on the manuscript.

